# Genetic properties underlying transcriptional variability across different perturbations

**DOI:** 10.1101/2024.04.15.589659

**Authors:** Saburo Tsuru, Chikara Furusawa

## Abstract

The rate and direction of phenotypic evolution depend on the availability of phenotypic variants induced genetically or environmentally. It is widely accepted that organisms do not display uniform phenotypic variation, with certain variants arising more frequently than others in response to genetic or environmental perturbations. Previous studies have suggested that gene regulatory networks channel both environmental and genetic influences. However, how the gene regulatory networks influence phenotypic variation remains unclear. To address this, we characterize transcriptional variations in *Escherichia coli* under environmental and genetic perturbations. Based on the current understanding of transcriptional regulatory networks, we identify genetic properties that explain gene-to-gene differences in transcriptional variation. Our findings highlight the role of gene regulatory networks in shaping the shared phenotypic variability across different perturbations.

## Introduction

The rate and direction of phenotypic evolution depend not only on natural selection but also on the availability of phenotypic variants, whether induced genetically or environmentally, upon which selection may act^1^. Even environmentally induced phenotypic variability or plasticity has the potential to initiate and direct adaptive evolution through processes such as genetic assimilation^2^. It is widely acknowledged that organisms exhibit heterogeneous phenotypic variation, with certain variants emerging more frequently than others in response to genetic or environmental perturbations^3, 4, 5, 6^. This bias in phenotypic variability can facilitate and constrain adaptive evolution across different phenotypic dimensions^7^, contingent on the alignment between the bias in phenotypic variability and the direction of selection^8^. Thus, without a thorough understanding of the nature of phenotypic variability, our understanding of why evolution has progressed in the manner that it does or why some phenotypes evolve repeatedly while others are one-offs remains incomplete.

Seemingly different types of phenotypic variability may actually be driven by a common mechanism. Theoretical studies have demonstrated that the bias in phenotypic variability is shared by different perturbations^9, 10, 11, 12, 13, 14^. Notably, several quantitative genetic studies identified similarity in the magnitude of variance, where traits sensitive to environmental perturbations tend to be sensitive to genetic perturbations^15, 16, 17, 18^. Additionally, there is a consistent pattern of directional variation among traits, with organisms frequently exhibiting correlated variation between traits^19^. Environmental influences on traits often act in the same direction as genetic influences, thereby shaping phenotypic variation in a consistent direction^6^. These similarities imply the existence of a common mechanism underlying the bias in phenotypic variability against any kinds of perturbations^12,13, 14, 20, 21^.

What molecular mechanisms bias the phenotypic variability? In biological systems, most traits arise from genetic interactions within networks represented by gene regulatory networks^22^. Genetic and environmental perturbations exert phenotypic effects via identical regulatory networks^23^. Consequently, phenotypic variance is expected to be influenced by the properties of the regulatory networks, regardless of the underlying causes of perturbations^8, 10^. Despite a growing number of studies demonstrating empirical evidence supporting the similarity both in susceptibility and in directionality of variations in gene expression levels between environmental and genetic perturbations^17, 18, 24^, it remains largely unknown how the real regulatory networks account for the observed bias in variability among genes^8^.

To address this issue, we characterize the variability of transcriptome profiles in *Escherichia coli* in response to different environmental and genetic perturbations. *E. coli* is an ideal model organism for this investigation because of its well-characterized gene regulatory network. Our analysis reveals that genes showing higher transcriptional variability against environmental perturbations tend to have higher sensitivity to genetic perturbations, indicating a shared gene-to-gene bias in variability across different types of perturbations. In addition, the major directions, or the top principal components, of correlated variation among thousands of genes are shared by different perturbations, demonstrating a common bias in the directionality of transcriptional changes. We identify 13 global transcriptional regulators that underlie this shared transcriptional variability across various perturbations. Genes regulated by these key regulators tend to exhibit greater transcriptional variability compared to those regulated by other transcriptional regulators. Moreover, these global regulators orchestrate coordinated transcriptional changes in their target genes, contributing to the predominant directionality of transcriptomic shifts observed across different perturbations. These empirical findings support the relevance of the gene regulatory networks in explaining the bias in phenotypic variability shared across different perturbations. Our results suggest that these biases are not random but are functionally significant in coping with fluctuating environments.

## Results & Discussion

### Similarity in transcriptional variance between environmental and genetic perturbations

We initially analyzed two public transcriptome datasets to evaluate the transcriptional variability across different perturbations. To this end, we curated transcriptome profiles from the PRECISE database^25, 26^, focusing on a single strain of *E. coli*, K12 MG1655, cultured under different environmental conditions (160 profiles, 76 unique environmental conditions) (**Supplementary Data 1, Supplementary** Fig. 1a,b), termed the Env dataset (**Fig. 1a**). Additionally, to evaluate transcriptional variability in response to genetic perturbations via natural evolution, we used transcriptome profiles from 16 natural strains of *E. coli* cultured under an identical environmental condition^27^, termed the Evo dataset (**Fig. 1b**). These isolates spanned the most common phylogroups and lifestyles of *E. coli*. Importantly, genetic mutations among these strains might reflect their diverse natural habitats, potentially confounding the estimation of transcriptional variability in response to genetic perturbations independent of environmental perturbations. To mitigate this, we conducted a mutation accumulation (MA) experiment^28^ using a mutator strain of *E. coli*, K12 MG1655, lacking the *dnaQ* gene^29, 30^ (**Fig. 1c**). Here, mutations primarily accumulate through genetic drift owing to small population bottlenecks, thus minimizing positive selection. We established two founder colonies (founders A and B) and propagated single colonies on lysogeny broth (LB) agar plates (yielding three MA lineages per founder). Consequently, we obtained transcriptome profiles from six MA lineages cultured in LB broth using the reliable DNA microarray technique^31^ (**Supplementary** Fig. 1c,d), termed the Mut dataset (**Supplementary Data 2**,**3**). Importantly, each dataset was generated within single laboratories using a common protocol, minimizing technical variation within each dataset. We used quantile-normalized and log2-transformed transcriptome data for the analysis.

**Fig. 1:**
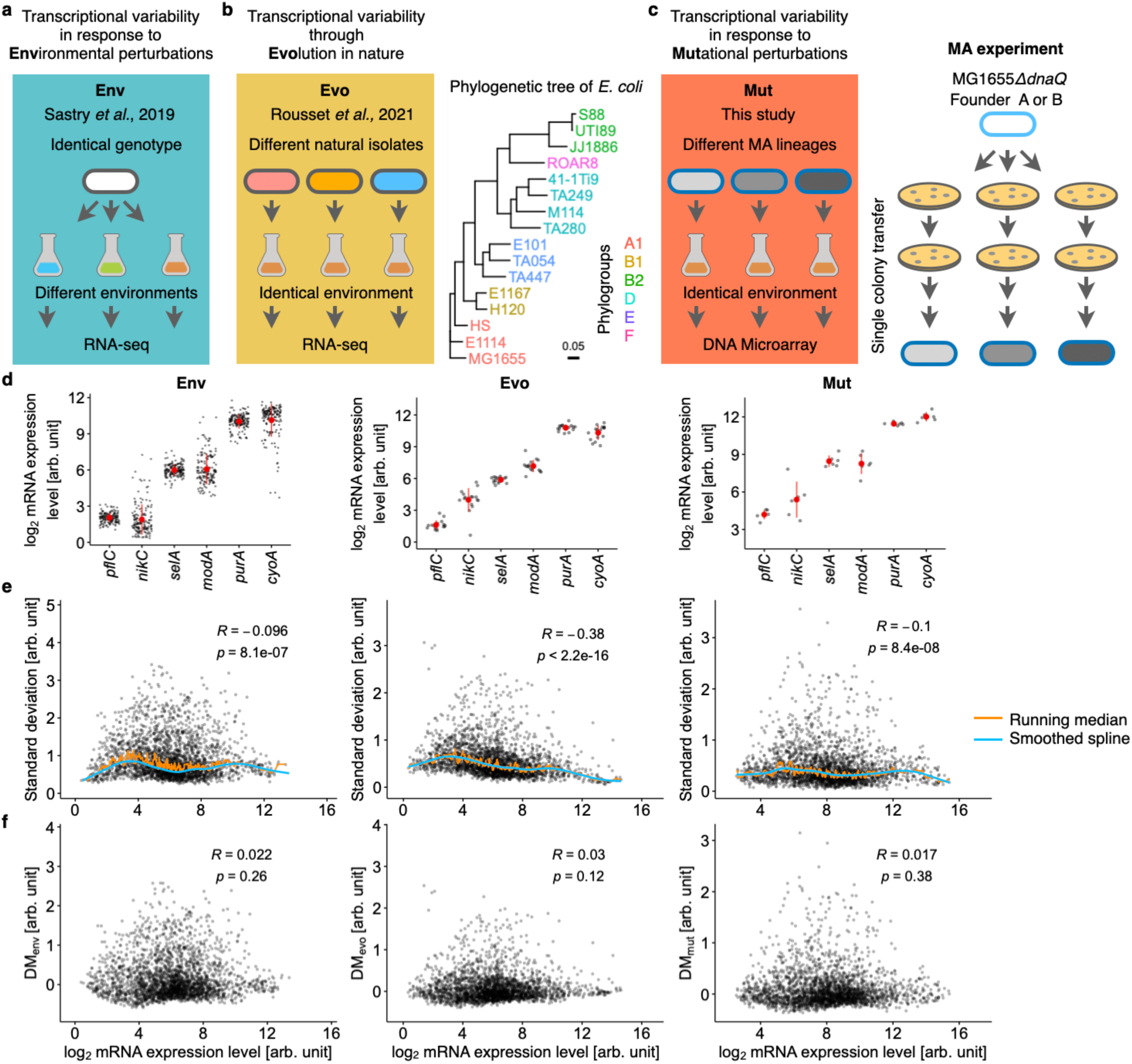
Transcriptional variability in *E. coli*. **a**, A schematic diagram of the Env dataset. The transcriptome profiles of *E. coli* MG1655 were obtained under various growth conditions^25, 26^. **b**, Schematic of the Evo dataset. The transcriptome profiles of 16 natural isolates of *E. coli* were obtained under a common growth condition^27^. The phylogenetic tree was based on the core genome alignment. **c**, Schematics of the MA and Mut dataset. The MA experiment was initiated by two founders of the mutator strain, *E. coli* MG1655*/idnaQ*. Single colonies were randomly picked and transferred daily to fresh LB agar plates for 30 rounds. Three MA lineages from rounds 10 and 30 for each founder were subjected to DNA microarray analysis. Transcriptome profiles were obtained during the exponential growth phase in LB. **d–f**, Transcriptional variability in Env, Evo, and Mut datasets (from left to right). The transcriptome profiles of the MA lineages in the final (30th) round were used for the Mut dataset. **d**, Expression levels of six representative genes. The expression levels were log2-transformed and quantile-normalized for each dataset. Each grey circle corresponds to a different bacterial culture. Large closed circles and error bars represent means and standard deviations for the transcriptome profiles of biologically independent samples: Env (n=160 profiles), Evo (n=16 profiles), and Mut (n=6 profiles). **e**, Relationship between means and standard deviations in the expression levels of 2622 genes present in all three datasets. Each dot corresponds to a different gene. The variability metric for each gene was defined as the vertical deviation from a smoothed spline calculated from the running median of the standard deviations. The calculated distance to the median is called DM, with subscripts of the corresponding datasets. For the Mut dataset, the standard deviations were obtained by calculating the variances in expression levels separately in each of the two groups distinguished by the founders (founders A and B) and subsequently averaging them. **f**, Relationship between DMs and mean expression levels. The insets represent Spearman’s rank correlation coefficients (R) and p-values (two-sided).

The means and standard deviations of expression levels were computed for each gene across each dataset (**Fig. 1d** for six representative genes). While standard deviation might directly measure transcriptional variability, it can be influenced by the mean^32^, leading to a confounding effect in estimating similarities in transcriptional variability between datasets. Notably, the standard deviations exhibited a statistically significant mean dependency for all datasets (**Fig. 1e**). To alleviate this influence, we calculated the vertical deviation of each standard deviation from a smoothed spline of the running median of the standard deviations (DM), which served as a measure of the magnitude of transcriptional variability. The DM values of each gene in the dataset was designated as DMenv, DMevo, DMmut for the Env, Evo, and Mut datasets, respectively (**Fig. 1f**, **Supplementary Data 4**).

Since each transcriptome profile for all datasets was obtained from millions of cells in unsynchronized cultures, gene expression levels are expected to be independent of the well-known temporal fluctuations in expression associated with the cell cycle. Unlike animals and plants, *E. coli* lacks developmental stages, and its temporal fluctuations in gene expression are primarily linked to replication-dependent changes in gene dosage ^33^. To verify that cell-cycle fluctuations did not confound our results, we examined the relationship between known temporal variability in mRNA expression and DM values. Pountain et al. performed single-cell RNA sequencing on the *E. coli* K12 MG1655 strain and quantified cell-cycle-dependent changes in gene expression. The amplitude of these fluctuations was measured as the fold change from trough to peak in mRNA levels within a cell cycle. Larger amplitudes represent higher temporal variability in mRNA expression without environmental or genetic perturbations. We confirmed that there was no positive correlation between DM values and the amplitude of cell cycle expression (**Supplementary** Fig. 2a). This is consistent with the expectation that our transcriptomic profiles reflect the population averages of unsynchronized *E. coli* cultures, and thus DM values did not reflect cell-cycle-associated temporal fluctuations.

In principle, replication-dependent changes in gene dosage could still influence DM values, even in unsynchronized cultures. At higher growth rates, overlapping cycles of replication occur, leading to higher copy numbers of genes located near the origin of replication (*oriC*). In slower growth conditions, this overlap does not occur, making gene dosage near *oriC* more sensitive to growth rate changes. This could potentially increase the transcriptional variability of genes near *oriC* across different environmental or genetic conditions. To test this, we explored the relationship between DM values and the genomic distance from *oriC* (**Supplementary** Fig. 2b). We found no significant negative correlation, suggesting that replication-dependent gene dosage effects did not significantly confound the DM values.

In addition to using unsynchronized cultures, we also accounted for the growth phase of the cultures in the Mut and Evo datasets. A previous study confirmed that transcriptome profiles obtained at different time points during the exponential growth phase under identical conditions are highly similar, especially when compared to profiles obtained under different environmental conditions or during other growth phases ^34^. To minimize the risk of unknown temporal fluctuations in expression levels in the absence of environmental changes, we performed transcriptomics on populations of exponentially growing cells under identical conditions for both the Mut and Evo datasets. Ideally, sampling the same cells multiple times under the same conditions could further validate these assumptions. However, both RNA sequencing and the microarray techniques used in this study are invasive methods for measuring gene expression, making this approach impractical. Unlike the other datasets, the Env dataset involved different growth conditions, and sampling times varied depending on the environment, meaning that some samples were taken outside the exponential growth phase. To assess the impact of these profiles on DMenv, we extracted the transcriptome profiles from the exponential growth phase within the Env dataset to construct a refined dataset, termed Envexp, based on information from the original papers in the PRECISE database. Recalculating DM values (DMenvexp) from this dataset (78 profiles, 37 unique environmental conditions, **Supplementary Data 1**), we observed a high correlation between DMenv and DMenvexp (**Supplementary** Fig. 3). Additionally, we confirmed that DMenvexp positively correlated with DMevo and DMmut, with correlation levels nearly equivalent to those of DMenv (**Fig. 2a**). These results indicate that samples taken outside the exponential growth phase did not significantly affect DMenv. Thus, we conclude that DMenv primarily reflects transcriptional changes in response to environmental perturbations, rather than temporal changes independent of environmental conditions. Overall, DM values are reasonably assumed to reflect transcriptional changes in response to environmental or genetic perturbations, rather than temporal fluctuations without such perturbations.

**Fig. 2:**
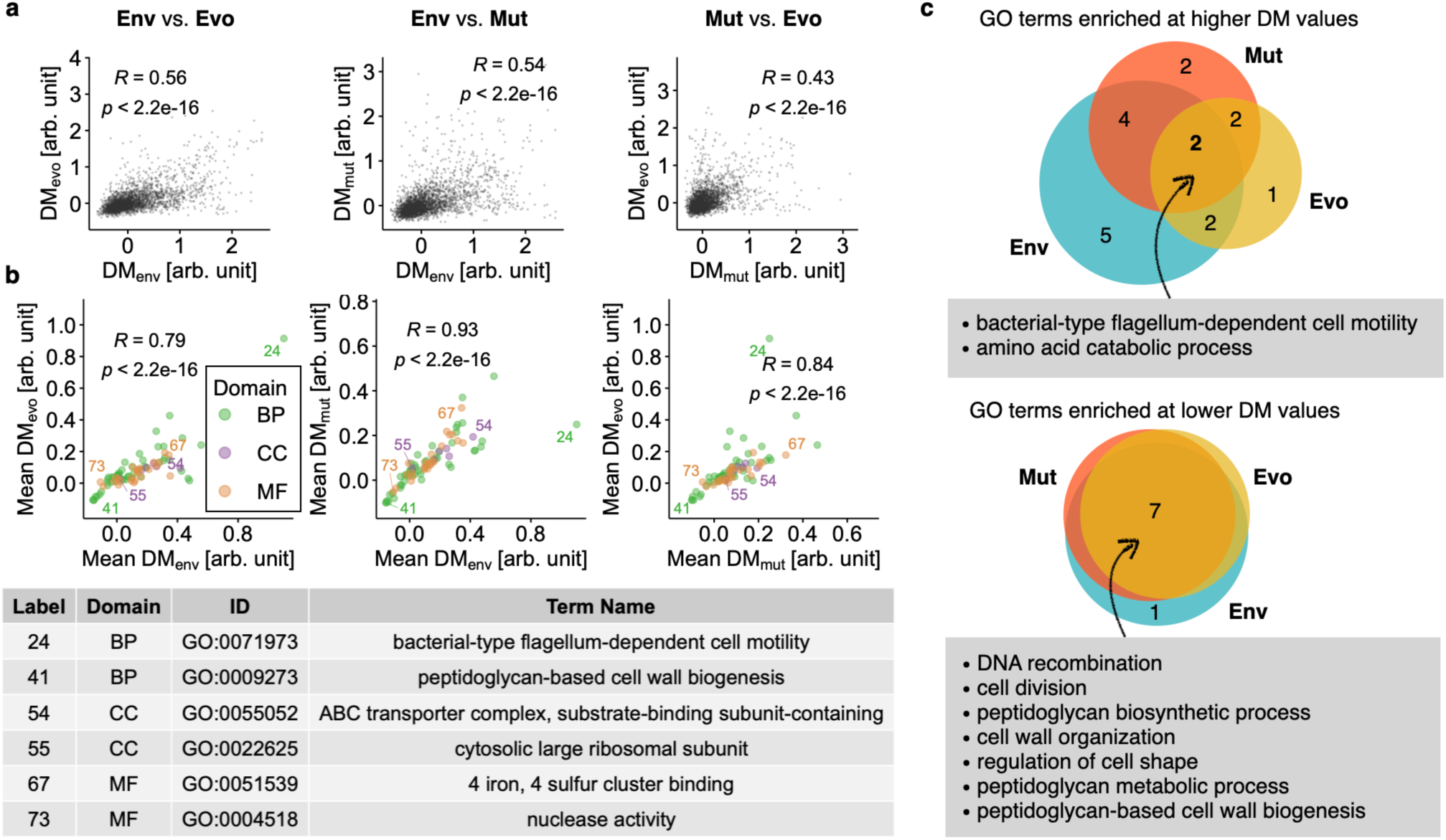
Positive correlations of transcriptional variability between datasets. **a**, Pairwise correlations are shown for DMs at the gene levels between the Env, Evo, and Mut datasets. **b**, Correlations of DMs at the functional category level between pairs of datasets. The genes were grouped into functional categories based on their GO terms. To reduce the redundancy of GO terms, a custom subset (82) of GO terms was used as detailed in the **Methods** section. Each dot corresponds to a different GO term. The mean DM was calculated by averaging the DM of the genes belonging to each GO term. The three domains of GO–biological processes (BP), cellular components (CC), and molecular functions (MF)–are labeled in different colors. Six representative GO terms are labeled with different numbers, as detailed at the bottom. Spearman’s R and p-values (two-sided) are shown in panels **a** and **b**. **c**, Venn diagram of GO terms enriched in higher (left) and lower (right) DM values in each dataset (GSEA, BH-adjusted p-values (q)<0.05, **Supplementary** Fig. 4). The Evo and Mut sets completely overlap in the right Venn diagram. The numbers indicated in the diagram show the number of GO terms enriched in the corresponding set.

Comparing DM values between datasets revealed positive correlations (Spearman’s rank correlation coefficient (R) =0.43–0.56, p<0.05) (**Fig. 2a**). These results indicate that genes with higher sensitivity to environmental perturbations tend to have higher sensitivity to genetic perturbations in terms of their expression levels. This phenomenon of canalization/decanalization could occur not only at the gene level but also at more complex phenotypes subject to natural selection. Canalization/decanalization in transcriptional changes has been observed in some functional categories of genes in yeast^18, 24^. For instance, genes involved in stress or amino acid metabolism tend to have higher transcriptional variability ^24^, while genes related to cell growth, maintenance, and cell cycles tend to have lower transcriptional variability against genetic perturbations, compared to random expectations ^18^. To explore whether functional associations in DM values were present in bacteria, we utilized a refined subset of Gene Ontology (GO) terms as functional categories (**Methods**) **(Supplementary Data 5**). Using the mean DM values of genes within each GO term as a measure of transcriptional variability, we observed positive correlations between the datasets even at the functional category level (**Fig. 2b**). To further investigate functional associations in DM values, we performed gene set enrichment analysis (GSEA^35^) with GO terms for both higher and lower DM values for each dataset independently. We found several GO terms significantly enriched in higher or lower DM values (BH-adjusted p-value (q)<0.05, GSEA). For instance, we identified genes related to 13 GOs that were significantly overrepresented at higher DM values compared with a uniform distribution. We identified several common GO terms across all datasets (**Fig. 2c**, **Supplementary** Fig. 4). For higher DM values, more than three GOs were found at every intersection between pairs of datasets. This overlap was significantly more than expected (p<0.0001) based on random drawing from a pool of 81 GOs without replacement. Notably, genes related to flagellum-dependent cell motility and amino acid catabolic processes were consistently enriched with higher DM values in all three datasets (q<0.05, GSEA). Tuning the activity of these functional categories in response to environmental changes is crucial for cells to cope with nutritionally fluctuating environments^36, 37^. In contrast, genes related to fundamental biological processes related to cell-wall construction and maintenance were consistently enriched at lower DM values for all datasets (q<0.05, GSEA, **Fig. 2c right**), indicating that the expression levels of genes within these categories were relatively robust to perturbations. The number of these common GOs (7) in lower DM values was significantly larger (p<0.0001) than expected (0.06) based on random drawing from a pool of GOs without replacement. We obtained relevant results using the functional categories defined by the KEGG pathways (**Supplementary** Fig. 5,6, **Supplementary Data 6**). There were positive correlations in DM values at the pathway level between the three datasets (Spearman’s R=0.5–0.67, p<0.05). Additionally, genes related to chemotaxes and peptidoglycan biosynthesis were overrepresented at higher and lower DM values, respectively, for all datasets (q<0.05, GSEA). These KEGG pathways were closely related to flagellum-dependent cell motility and cell-wall construction in GO terms, respectively. These results indicate frequent associations between canalization/decanalization in transcription level of genes and their functions, suggesting a certain common mechanism underlying the heterogeneous transcriptional variability associated with functional categories.

### Transcriptional regulators influencing transcriptional variability

To explore the underlying mechanism, we examined the influence of transcriptional regulators on transcriptional variability. First, we conducted GSEA for each dataset independently based on known regulatory interactions (**Supplementary Data 7**) to identify transcriptional regulators that affect target genes exhibiting higher and lower DM values, termed TRhigh and TRlow, respectively. We analyzed two golden standard databases, RegulonDB^38^ and EcoCyc^39^, collecting known regulatory interactions between genes. We focused on the transcription factors and sigma factors that regulate more than four and less than 1001 target genes. For the Env dataset, we identified 30 transcriptional regulators enriched in higher DMenv, such as FlhDC and Fur, and termed them TRenv (q<0.05, GSEA, **Fig. 3a**). This enrichment means that the target genes of these regulators were overrepresented at higher DMenv compared with a uniform distribution. In contrast, we identified only four transcriptional regulators enriched in the lower DMenv (q<0.05, GSEA). Similar analyses were performed for DMevo and DMmut, independently (**Supplementary** Fig. 7). Consequently, we identified 29 (TRevo) and three transcriptional regulators enriched in higher and lower DMevo, respectively. Similarly, 28 (TRmut) and two transcriptional regulators were enriched in the higher and lower DMmut, respectively (q<0.05, GSEA). Thus, fewer transcriptional regulators were overrepresented at lower DM values, suggesting that regulation by transcriptional regulators tends to increase the transcriptional variability of target genes, with some regulatory motifs, such as negative feedback, potentially repressing fluctuations in gene expression, as a minor exception^40^.

**Fig. 3:**
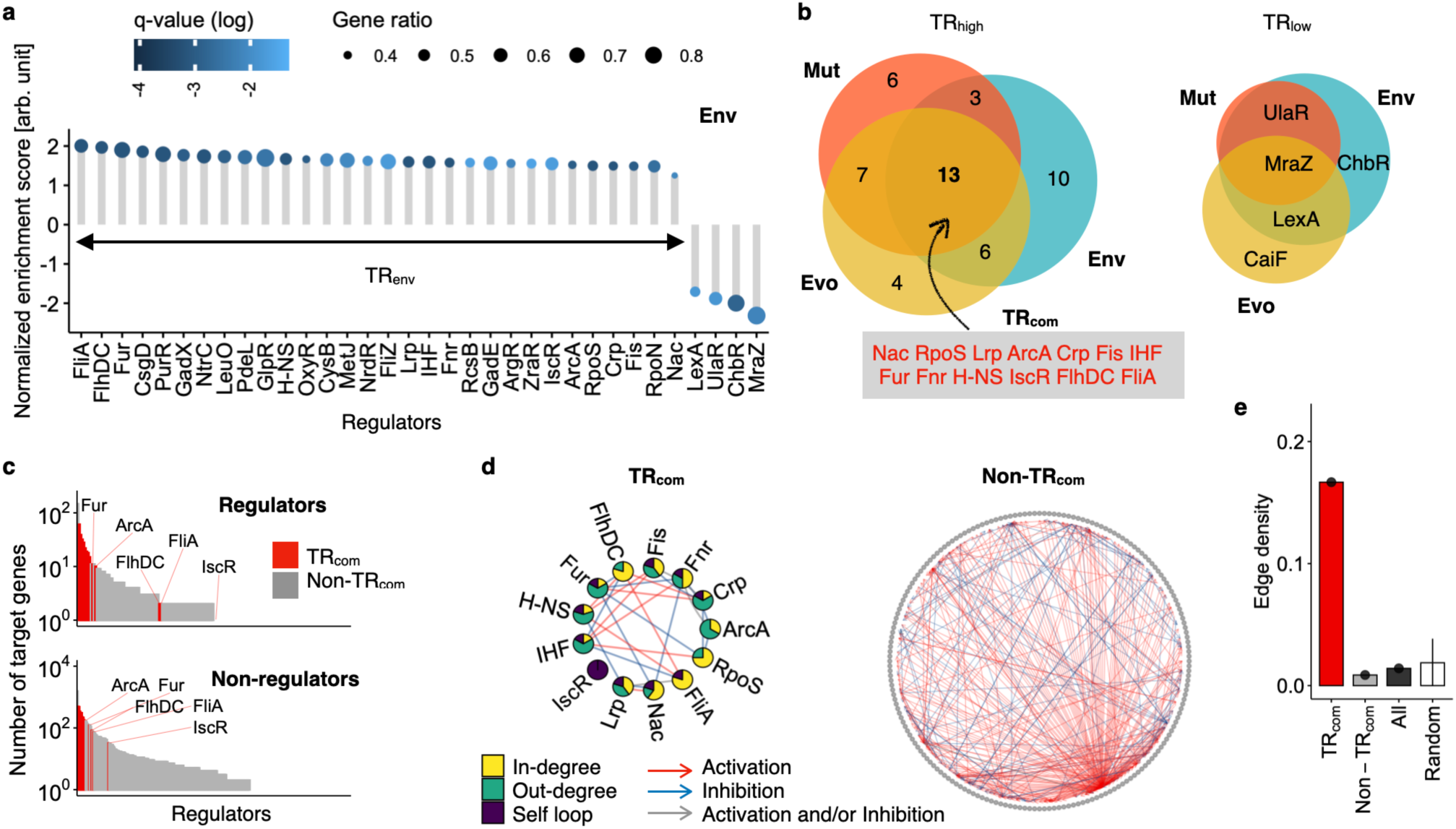
Transcriptional regulators and transcriptional variability. **a**, Transcriptional regulators (TR) enriched in genes with higher and lower DMenv. GSEA identified statistically significant TRs (q<0.05). Positive and negative enrichment scores indicate enrichment at higher and lower DMenv, respectively. The gene ratio was calculated as the number of core target genes contributing to enrichment scores divided by the number of target genes. TRs with positive enrichment scores are labeled as TRenv. The q-values (adjusted p-values) were computed using the BH procedure. **b**, Venn diagram of the enriched TRs in each dataset. TRevo and TRmut represent the TRs with positive enrichment scores in the Evo and Mut datasets, respectively. The intersection of the three sets represents TRcom. **c**, Rank plot of the number of target genes in the TRs. The TRs were sorted in decreasing order. Target genes were divided into regulators (transcription factors and sigma factors, upper panel) and non-regulators (lower panel). The bars are colored to indicate whether the TRs belong to TRcoms or Non-TRcoms, with . labels representing subsets of TRcoms. **d**, Circular representation of *E. coli* transcriptional regulatory network (TRN) within TRcoms (left) and Non-TRcoms (right). Edges are colored based on the type of regulation (activation, inhibition, and activation and/or inhibition). Pie charts indicate the percentage of each degree within the TRcom nodes. The edges corresponding to the self-loops were omitted from the visualization. **e**, Edge densities within regulatory groups. The random group was generated by drawing 13 TRs at random 10,000 times from a pool of TRs, where TRs that regulate fewer than five target genes were omitted. The bar and error bar for the random group represent the mean and standard deviation (n=10,000), respectively. By definition, the sample size for the other groups was one. Edge density was calculated as the total number of edges within the regulators divided by the total number of possible edges within the regulators, excluding the self-loops.

Next, we explored the similarity between TRenv, TRevo, and TRmut, which were identified in each dataset independently. We found that 13 transcriptional regulators, referred to as TRcom, were common among the three datasets, corresponding to more than one-third of the transcriptional regulators enriched in each dataset (**Fig. 3b**). The number of TRcoms (13) was significantly larger (p<0.0001) than the expected number (1.7) for the intersection of the three sets, constructed by random drawing from a pool of 118 TRs without replacement. These results suggest that the target genes regulated by TRcom exhibited higher transcriptional variability in response to both environmental and genetic perturbations, indicating that TRcoms function as a common mechanism to bias transcriptional variability against various perturbations.

### Network signatures of transcriptional regulators influencing transcriptional variability

How do TRcoms enhance transcriptional variability of target genes against a variety of perturbations both inside and outside of the cells? We found that TRcoms are global regulators with a large number of target genes, including TRcoms themselves (**Fig. 3c**). Based on these findings, we speculated that TRcoms might regulate each other frequently, which could increase the probability of the propagation of different perturbations to their target genes. To test this hypothesis, we explored the regulatory structure of a network comprising TRcoms alone (**Fig. 3d**). We found that most TRcoms had both an in-degree and an out-degree (11 out of 13 regulators). FliA had no out-degrees but four in-degrees. Only IscR was isolated with no regulatory interactions within TRcoms. This network structure indicates that the network within TRcoms is not top-down structured but has many mutual interactions (activations and inhibitions) within TRcoms. The edge density within the TRcoms (**Fig. 3e**) was remarkably high (0.17), with statistical significance (p<0.0001), compared to the edge density of random expectations (0.019). These results indicate that the TRcoms form a relatively densely connected network that is proficient in propagating the influence of perturbations within this group.

Next, we explored the effects of TRcoms on transcriptional variability. First, we compared the DM values of the target genes with TRcom-containing regulation with those of the target genes with TRcom-less regulation (**Fig. 4a**). The target genes of TRcoms tended to have higher DM values in all datasets than other genes with the same number of unique regulators. We found none or very weak positive correlations between the DM values of the genes without regulations by TRcoms and the number of unique regulators that the genes received in all datasets (Spearman’s R=∼0.13). On the other hand, the target genes of TRcoms exhibited relatively high positive correlations in all datasets (Spearman’s R=0.23–0.35, p<0.05). These results indicate that the recruitment of TRcoms is more effective in enhancing transcriptional variability. To further investigate this hypothesis, we explored the relationship between DM values and the number of unique TRcoms among the genes with a fixed number of regulators. In the case of the genes that received five unique regulators (**Fig. 4b**), we found a positive correlation between DMenv and the number of unique TRcoms among the five regulators (Spearman’s R=0.29, p<0.05). We performed similar analyses for all the datasets and found pervasive positive correlations across broad classes of the number of unique regulators (**Fig. 4c**). These results highlight the role of TRcoms in enhancing transcriptional variability through dense mutual regulation within TRcoms and co-regulation by multiple TRcoms, likely because both increase *trans*-target sizes^18^ that affect the expression levels of the genes regulated by TRcoms in response to both environmental and genetic perturbations.

**Fig. 4:**
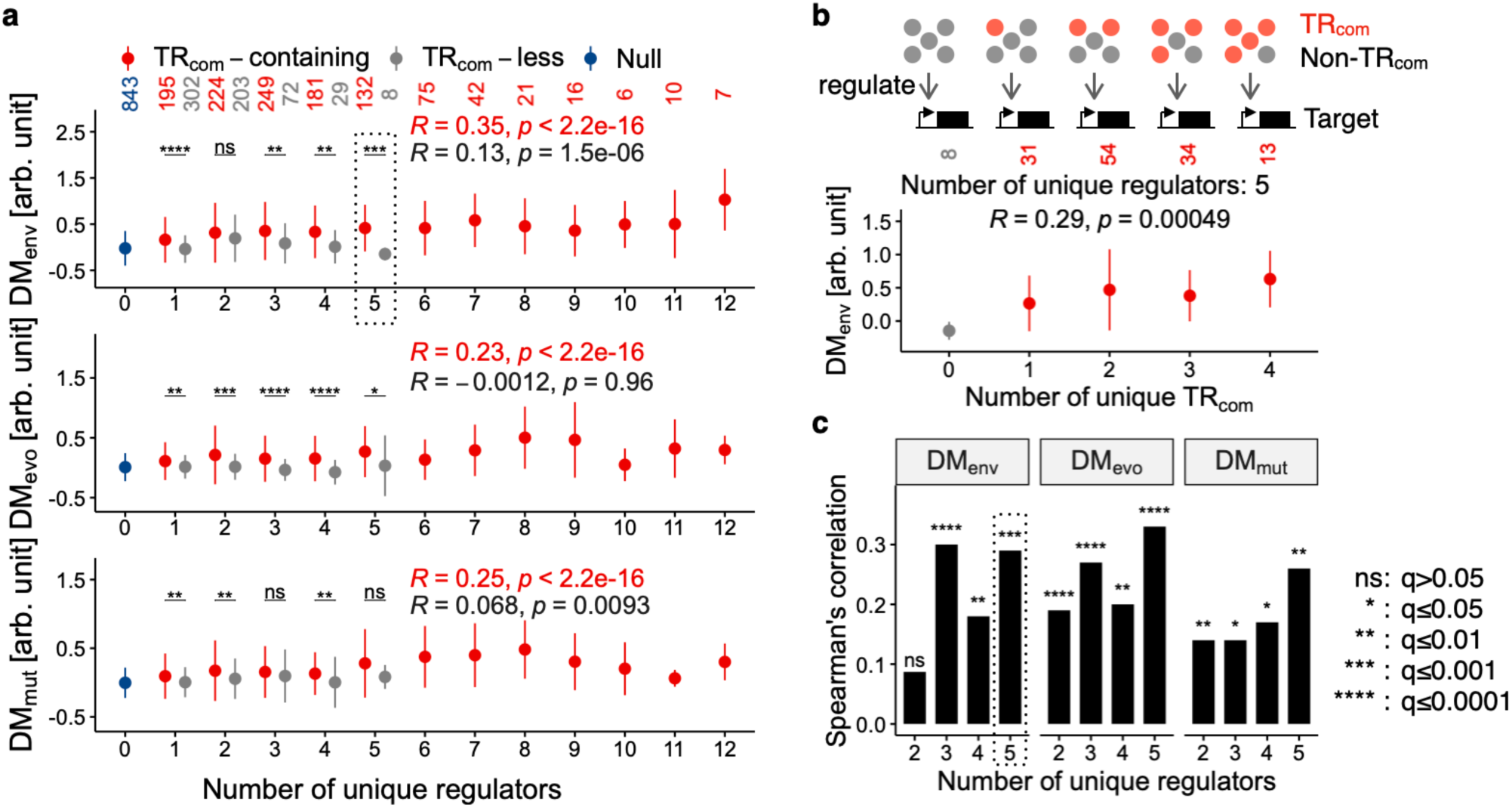
Relationship between transcriptional variability and the number of transcriptional regulators. **a**, Relationship between DM values and the number of unique regulators. The top, middle, and bottom panels show the DMenv, DMevo and DMmut, respectively. Target genes receiving TRcom-containing regulation, TRcom-less regulation, and genes without known regulators are labeled in different colors. Means and standard deviations are shown as the points and the error bars, respectively. The sample sizes (n) are indicated at the top. Data points for TRcom-containing and TRcom-less regulations were restricted to those comprising more than four target genes. Spearman’s R and p-values (two-sided) were calculated for data points 0–5 on the x-axis for both TRcom-containing and TRcom-less regulations. Asterisks represent the statistical significance levels (q-values) of the Wilcoxon test between TRcom- containing and TRcom-less genes in each class. The target genes within the dotted square are used in panel **b**. **b**,**c**, The correlation between DM values and the number of TRcom regulations. **b**, Relationship between DMenv and the number of TRcoms for target genes regulated by any of the five regulators. Means and standard deviations are shown as the points and the error bars, respectively. The sample sizes (n) are indicated at the top. Spearman’s R and p-values (two-sided) are shown. **c**, Spearman’s R for DMenv (left), DMevo (center), and DMmut (right). The enclosed square corresponds to panel **b**.

To understand how TRcoms access a variety of perturbations, we explored the relationship between TRcoms and the known molecular machinery for sensing stimuli both inside and outside the cells. First, we performed bow-tie decomposition of the TRN to identify the subnetworks to which the TRcoms belonged (**Fig. 5a**–**d**, **Supplementary Data 8**). The *E. coli* TRN was formally dissected into seven sectors defined by Yang’s bow-tie structure^41^ based on the directed regulatory links between genes^42^. The primary sectors are IN, OUT and the greatest Strongly Connected Component (SCC). The SCC sector represents the subnetwork, in which each node (TR) is reachable by any other node following the direction of regulation from the TR to its target. The IN sector comprises TRs that are not included in the SCC and can reach TRs in the SCC. The OUT sector comprises nodes (TRs or non-regulatory genes) not included in SCC but reachable by TRs in SCC. The numbers of nodes in these primary sectors in *E. coli* TRN were seven TRs, 62 TRs and 2715 genes including the TRs for IN, SCC, and OUT, respectively (**Fig. 5e**). As expected from their many target genes, TRcoms (12 of 13 TRcoms) and TRhighs (26 of 48 TRhighs in Env, Evo, and Mut) belonged to SCC (**Fig. 5d**). This implies that TRcoms can receive broad perturbations from approximately 70 genes in the IN and SCC sectors. We noted that TRcoms in SCC still showed a higher edge density (0.19) than other TRs in SCC (p<0.005), supporting the especially dense connections within the TRcoms.

**Fig. 5:**
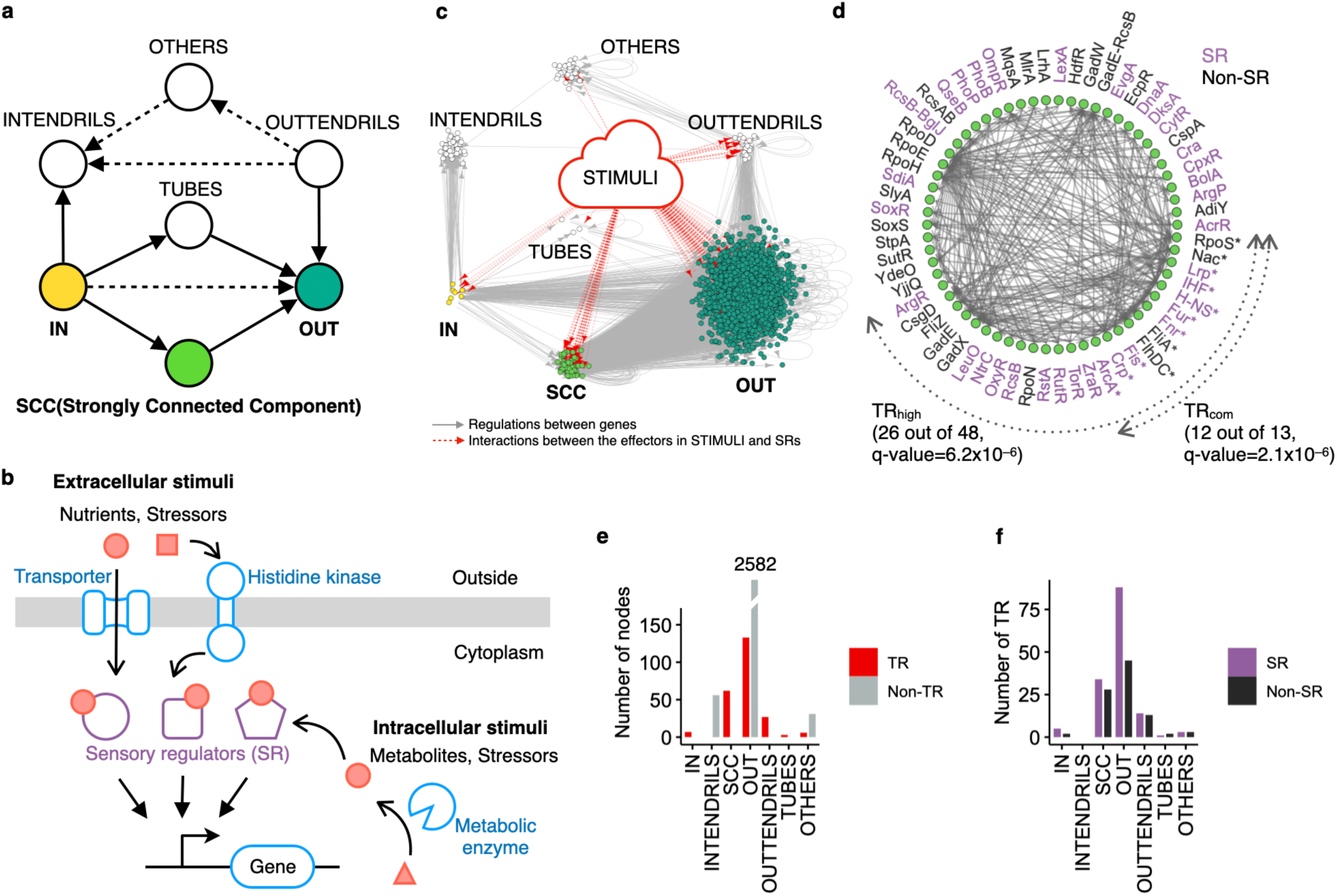
Bow-tie decomposition of the gene regulatory network in *E. coli*. **a**, Yang’s bow-tie structure^41^ is delineated into seven sectors, with the main sectors highlighted: IN, OUT, and the strongly connected component (SCC). Arrows indicate the direction of regulation, with solid arrows defining each sector and dashed arrows representing possible regulations without violating the sector definitions. **b**, Schematic illustrating how sensory regulators (SR) influence the expression levels of target genes by interacting with intracellular and extracellular effectors (nutrients, metabolites, and stressors). **c**, Bow-tie representation of the gene regulatory network in *E. coli* based on panel **a**. Each node represents a TR or target gene. Solid arrows denote regulation (activation and/or inhibition). The STIMULI block represents an aggregated group of known effectors interacting with SRs, with dotted arrows indicating the interactions between the effectors in STIMULI and SRs. **d**, Regulatory network within the SCC. Each node represents a TR. SR and non-SR groups are labeled in different colors, with asterisks representing TRcom. **e**, Distribution of nodes in each sector. TR and non- TR groups are labeled in different colors. IN, SCC, TUBES, and OUTTENDRILS consist solely of TR, whereas INTENDRILS contains only non-TRs by definition. **f**, Number of TR in each sector. SR and non-SR groups are labeled in different colors.

Next, we investigated how TRcoms sense different intracellular and extracellular stimuli. *E. coli* employs at least 145 sensory regulators to transmit various stimuli to the TRN (**Fig. 5b**, **Supplementary Data 8**). These regulators, termed SRs, change their regulatory activity depending on their interactions with corresponding effectors, as summarized by Martínez-Antonio *et al*^43^. For visualization, we defined the STIMULI block as an aggregated group of the known effectors that interact with SRs, such as nutrients, stressors and metabolites. We found that the IN and SCC sectors contained 39 SRs (27% of the total SRs, **Fig. 5f**), which was reasonable agreement with the random expectation based on the number of TRs in these sectors. Interestingly, more than half of TRcoms (8 of 13) were SRs. These results indicate that TRs in the SCC sectors, TRcoms included, not only serve as intermediate managers in information processing but also play a role in sensing different kinds of stimuli. These characteristics of TRcoms, particularly their presence in SCC with many SRs, may enable TRcoms to access a variety of stimuli both inside and outside the cells.

### Similarity in direction of transcriptional variance between environmental and genetic perturbations

Next, we explored the major directions of correlated transcriptional variation among thousands of genes. Theoretically, genetically caused phenotypic variance is expected to be the greatest along the major axis or the principal component of environmentally caused phenotypic variance^10, 12, 13^. To identify such principal components (PC) and test this prediction, we performed principal component analysis (PCA) for each dataset (**Fig. 6a**, **Supplementary** Figs. 8–10). The first three principal components (PC1–3) explained 46%, 53%, and 57% of the total variance in the Env, Evo, and Mut datasets, respectively, indicating their significance as the major axes of transcriptional variation for each dataset. Correlation analysis based on the loadings of these major axes revealed similarities in their directions across the datasets (**Fig. 6b**). We used the absolute values of Spearman’s R because of the relative nature of the signs of the principal components. Specifically, PC1 in the Env dataset exhibited relatively strong correlations with PC1 and PC2 in the Evo dataset (Spearman’s R = 0.45 and 0.41, respectively, p<0.05) and with PC1 and PC2 in the Mut dataset (Spearman’s R = 0.34 and 0.41, respectively, p<0.05). Similarly, PC1 in the Evo dataset showed a relatively strong correlations with PC1–3 in the Env and Mut datasets. These correlations in loadings were much stronger than expected (∼0.03 in Spearman’s R) based on independent datasets comprising no correlations in expression levels between genes (**Supplementary** Fig. 11c). To validate the high similarity of the PCs between the three datasets more formally, we assessed the extent to which the principal components in the Env dataset explained the transcriptional variance in the other two datasets. Overall, all principal components in the Env dataset explained up to 37% and 39% of the total variance in the Evo and Mut datasets, respectively (**Fig. 6c**). These fractions were much larger than random expectations (∼3%, **Supplementary** Fig. 11d). Moreover, the cumulative variance plot displayed a convex upward curve for both the Evo and Mut datasets, as well as Env, which was distinct from a linear-like curve in random expectations (blue line in **Supplementary** Fig. 11d). These findings support the notion that the major directions of transcriptional variation in response to environmental perturbations are considerably similar to those observed in response to genetic perturbations, despite a large number of genes.

**Fig. 6:**
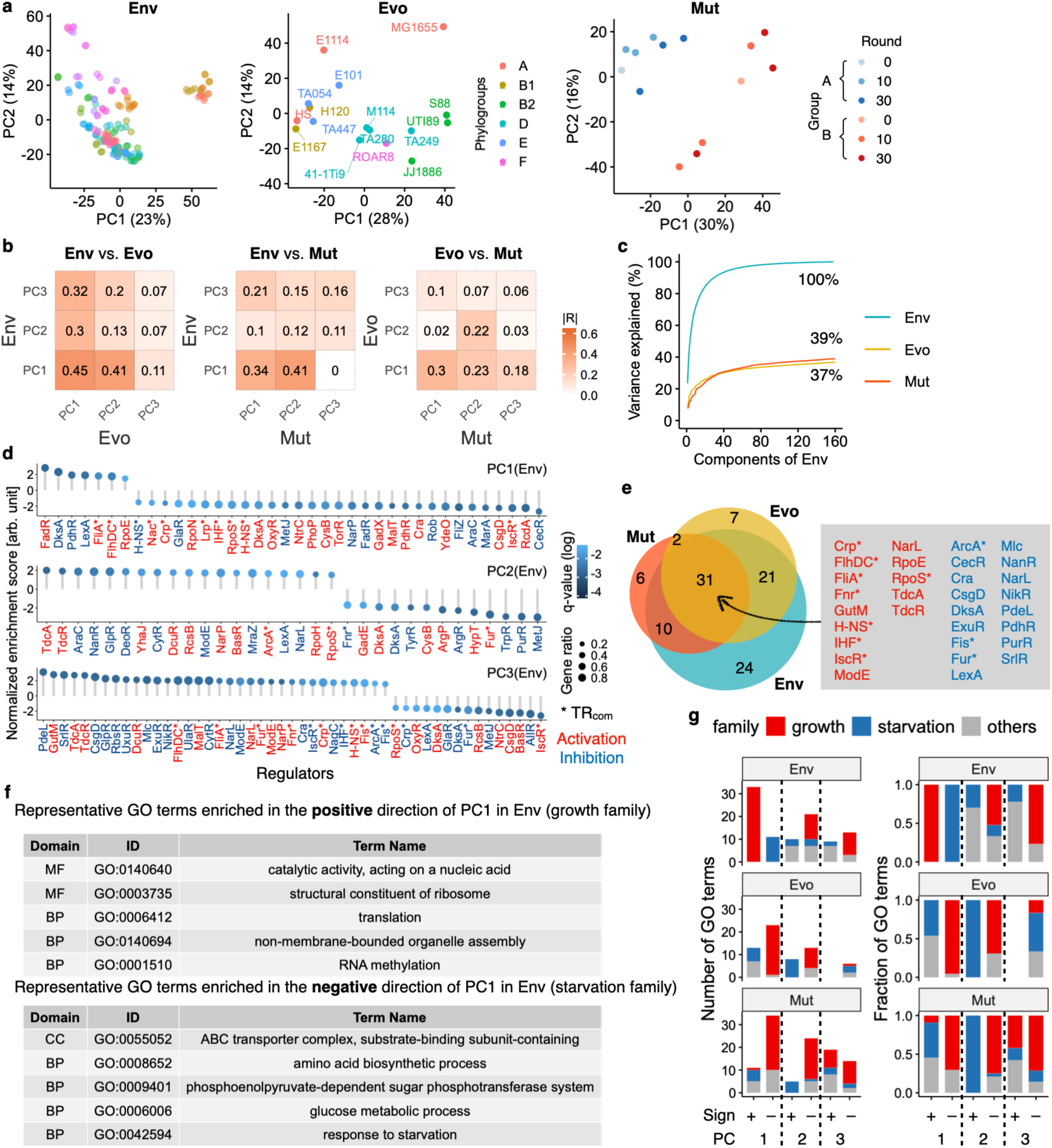
Alignment of genetic and environmental correlations in transcriptional variability. **a**, First and second principal components (PC) of Env, Evo and Mut arranged from left to right. Points are colored by environmental condition in Env. **b**, Correlation analysis for loadings of the first three PCs among the three datasets. The color scale and numbers in the tiles represent the absolute values of Spearman’s R. **c**, Cumulative variance explained by the PCs defined by the Env dataset. Overall, the Env dataset explained 37% and 39% of the variance in the Evo and Mut datasets, respectively. **d**, TRs were significantly enriched (q<0.05) in greater loadings, either positive or negative, of the first three PCs in the Env dataset. GSEA was used to identify regulations, distinguishing between activatory and inhibitory regulation. Asterisks denote regulations by TRcoms. **e**, Venn diagram illustrating the TRs enriched in the first three PCs of each dataset. GSEA was performed for each dataset, as shown in panel **d**. Numbers indicate the number of regulations in the corresponding set. The detailed regulations at the intersection of the three sets are shown on the right. **f**, GO terms enriched with greater loadings in the positive (top) and negative (bottom) directions in PC1 of the Env dataset (five representatives). The former and latter GO terms were categorized into the growth and starvation families. **g**, Number (left) and fraction (right) of GO terms enriched in each sign of each PC in each dataset. GSEA was conducted for each PC in each dataset. The fraction was calculated by dividing the number of GO terms in each family by the total number of GO terms in each PC sign. A full list of the enriched GO terms is presented in **Supplementary** Fig. 14–16.

A directional bias in transcriptional variations is expected to reflect transcriptional regulations^8^. To validate this prediction, we explored the transcriptional regulators that contribute to the major directions. Utilizing the loadings of PC1–3 in each dataset, we conducted GSEA (**Fig. 6d**,**e**). We found the transcriptional regulations enriched in the positive and negative directions of PC1–3 in the Env dataset (**Fig. 6d**). For instance, activation by FadR and FlhDC or inhibition by DksA and PdhR was enriched in the positive direction of PC1. Conversely, activation by RcdA and IscR or inhibition by CecR and MarA was enriched in the negative direction of PC1. We conducted a similar GSEA for the other two datasets (**Supplementary** Fig. 12), identifying 31 regulations common across all three datasets (**Fig. 6e**), including 11 regulations by TRcoms, which is consistent with the notion that TRcoms are global regulators collectively controlling numerous targets. The number of common regulations (31) was significantly larger (p<0.0001) than the expected number (0.2) for the intersection of the three sets constructed by random drawing from a pool of 267 regulations without replacement. Overall, these findings support the idea that the similarity in the direction of transcriptional changes in response to different types of perturbations arises from shared transcriptional regulation. To further confirm the relevance of these findings, we analyzed transcriptome profiles of *E. coli* mutants harboring rewired transcriptional regulations^44^ that perturb native transcriptional regulations between major global regulators (**Supplementary** Fig. 13). Each of the rewired strains has a unique shuffled synthetic operon comprising a plasmid copy of the native promoter region of a global regulator (‘A’) and a plasmid copy of a gene coding for another global regulator (‘B’). Accordingly, promoter-mediated transcriptional regulations controlling expression of ‘A’ are newly connected to ‘B’ without mutating native chromosomal regulations. Consequently, these rewired strains have new regulatory “links” between randomly selected pairs of global regulators, perturbing native transcriptional regulations between global regulators. The transcriptome profiles of the rewired strains were originally obtained by Baumstark *et al*^45^ under identical environmental conditions. Using this dataset, termed the Rwr dataset, we performed PCA and explored how the directionality in transcriptional variance differed from the common directionality shared by the three main datasets (**Supplementary** Fig. 13c,d). We confirmed that the top two PCs in Rwr showed lower correlations (∼0.18 at most) with those in the three main datasets, which was very different from the higher correlations between the three main datasets (**Fig. 6b**). Relatively higher correlations were found between the third PC in Rwr and the top two PCs in the three main datasets. Consistently, the PCs defined by the Env dataset explained at most 29% of the variance in the Rwr dataset, which was lower than those of the Mut and Evo datasets (**Supplementary** Fig. 13e). These results indicated that the directionality in transcriptional variation between rewired strains was less similar to the common directionality shared in the other datasets. These results highlighted the importance of regulatory networks between global regulators in major directions in transcriptional variability, supporting the relevance of our findings.

Furthermore, functional enrichment in GO terms was observed in the top three PCs across all three datasets (**Fig. 6f**, **Supplementary** Figs. 14–16). The most major direction in Env (PC1) revealed enrichment of translation, cell division, and cell-wall-related biological processes in one direction (positive sign), and responses to starvation and amino acid catabolic/biosynthetic processes in the opposite direction (negative sign). We termed these opposing functional families growth and starvation families, respectively, because of their close association with cellular growth and starvation states in response to environmental conditions^46^. The correlation between PC1 in Env and PC1 and PC2 in Evo and Mut suggested that approximately half to one-thirds of the functional categories in both families were enriched in these PCs, with opposite directions between the families retained across datasets. Importantly, the mutually opposite directionality observed in these functional families tended to be preserved across different perturbations, suggesting a consistent response pattern at the functional category level (**Fig. 6g**).

### Similarity of phenotypic variability across various perturbations

Heterogeneous phenotypic variability is not a random occurrence devoid of biological significance (**Fig. 2c** and **6f,g**), but rather it may result from or be facilitated by the adaptive generalization of past environments^9, 10, 11, 12, 13^. In addition, several studies suggest that evolution tends to produce global canalization that buffers phenotypes against not only genetic and environmental perturbations^12, 13, 21^, but also perturbations caused by stochastic molecular noise^47, 48^. These findings imply a similarity in variability in gene expression levels across a broader range of perturbations than previously examined. For instance, genes with higher/lower DM values in Env might also have higher/lower transcriptional variability in response to other environmental perturbations or other nongenetic perturbations (molecular noises) untested in the Env dataset. Similarly, genes with higher/lower DM values in response to random mutations accumulated through genetic drift (Mut) might exhibit higher/lower DM values in response to beneficial mutations accumulated by strong selection in adaptive evolution. To address these possibilities and validate a broader applicability of the findings from the main datasets, we analyzed five additional datasets for *E. coli*, termed Strs_hp, Strs_he, Strs, Prtn, and Noise_ch (**Fig. 7a**, **Supplementary** Figs. 17–24, **Supplementary Note 1**). These datasets include transcriptome or proteome profiles obtained under relatively narrower or more specific conditions than the three main datasets (Env, Evo, and Mut), but under several perturbations absent in the three main datasets, except for Prtn. The Prtn dataset, which contains several environmental conditions similar to those used in Env, was used to test whether the bias in transcriptional variability was retained at the protein level.

**Fig. 7:**
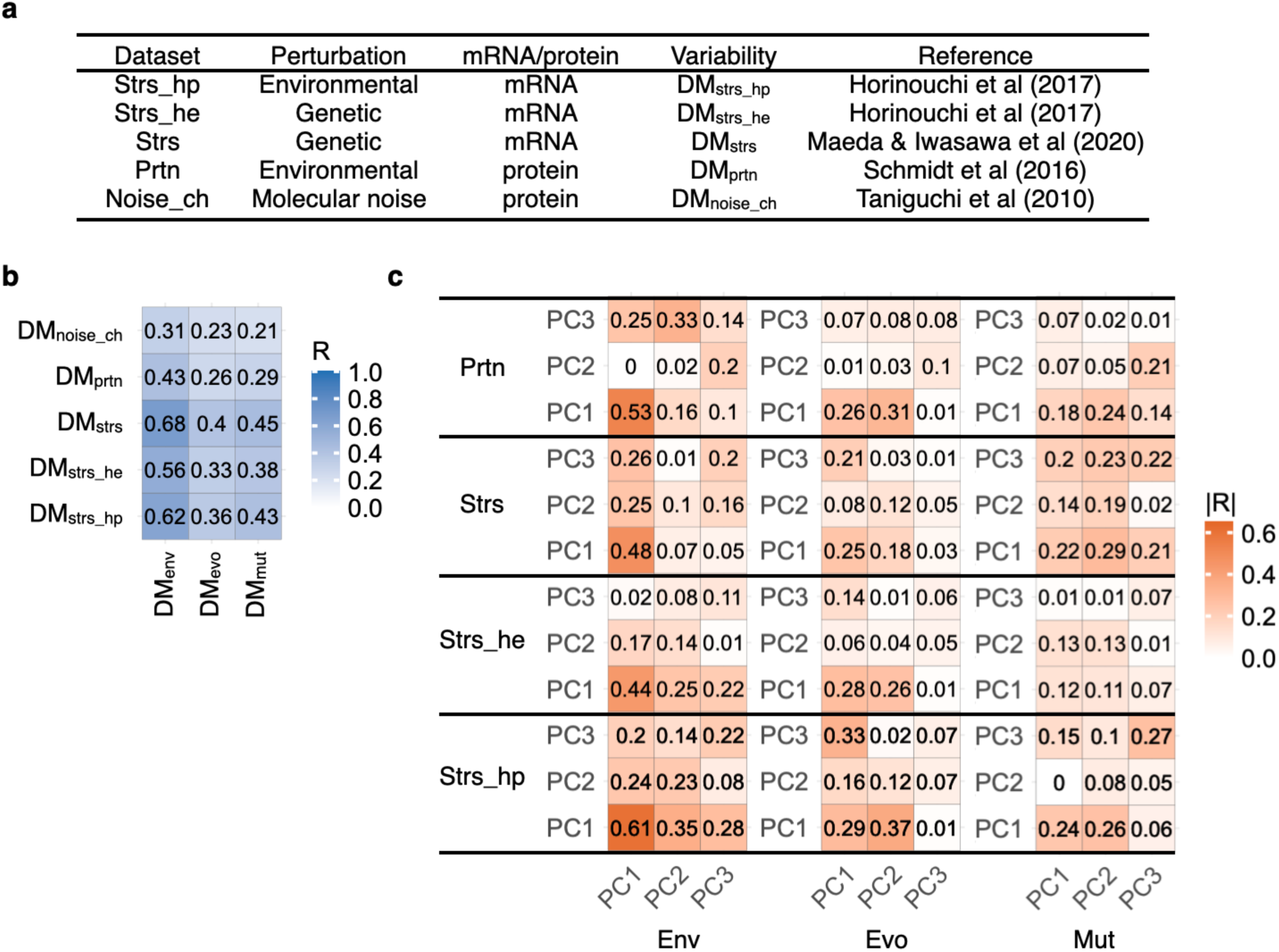
Similarity of phenotypic variability across different datasets. **a**, Datasets used to compare similarity in both variability (**b**) and directionality (**c**) of gene expression levels (mRNA/protein) with three main datasets. **b**, Correlation analysis of the variability across different datasets. The color bar and numbers in the tiles represent Spearman’s R. Correlation analysis for DMnoise_ch, was conducted only for highly expressed proteins with extrinsic noise dominance. The results of the lowly expressed proteins are detailed in **Supplementary Note 1**. **c**, Correlation analysis of loadings of the first three PCs between the datasets. The color bar and numbers in the tiles represent absolute values of Spearman’s R.

The phenotypic variabilities based on these datasets, either at the transcriptional or protein level, were termed DMstrs_hp, DMstrs_he, DMstrs, and DMnoise_ch as detailed in **Supplementary Note 1**. In short, DMstrs_hp and DMprtn show variability in their expression levels (mRNA for Strs_hp and protein for Prtn) in response to various environmental perturbations. DMstrs_he and DMstrs represent transcriptional variability in response to genetic mutations accumulated through adaptive laboratory evolution under various stressors. DMnoise_ch represents the protein noise of clonal populations under identical environmental and genetic conditions. We found that these DM values were positively correlated with those of the three main datasets (**Fig. 7b**). Notably, the correlations with DMenv show relatively strong in all pairs (Spearman’s R = 0.31–0.68, p<0.05). Next, we conducted PCA to identify the directionality of each dataset. To understand the similarity in the major directions across the datasets, we compared PC1–3 of these additional datasets with those of the three main datasets (**Fig. 7c**). Correlation analysis using the loadings of PC1–3 revealed that PC1s of the additional datasets tended to correlate with PC1s or PC2s of the three main datasets. In particular, PC1 in the Env dataset showed the strongest correlation with the PC1s of the additional datasets. Thus, transcriptional variability tended to be retained at the protein level, as shown in the Prtn dataset, suggesting the impact of transcriptional regulation on the variability in phenotypes governed by proteins. The similarity in transcriptional variability with Strs_hp supported the impact of the transcriptional variability identified in the three main datasets on broad environmental conditions. Importantly, the similarity with Strs and Strs_he highlighted the impact of transcriptional variability under relaxed selection (Env or Mut) on the evolutionary diversification in transcriptional phenotypes accompanied by adaptive evolution. Interestingly, the transcriptional variability in response to mutations (Mut) was also coupled with stochastic protein noise (Noise_ch), which was previously reported in yeast ^18^. Taken together, these results indicated similar trends in gene-to-gene differences in both the magnitude and major directions of expressional variations across various perturbations, suggesting global canalization/decanalization across different perturbations^12, 13, 14, 21, 47^.

### Concluding remarks

In this study, we characterized the transcriptional variability of *E. coli* against different perturbations and explored the regulatory commonalities between genetic and environmental perturbations. We identified common global transcriptional regulators that underlie the shared variability across different perturbations. These transcriptional regulators interact relatively densely with each other and strongly connect with many sensory regulators, forming a network structure that is favorable to facilitate the frequency of propagation of various stimuli both inside and outside the cells. In addition, the more regulated genes by these influential regulators showed greater changes in expression effectively. PCA revealed that these global transcriptional regulators simultaneously induced changes in the expression levels of multiple genes and contributed significantly to the formation of major directions of genome-wide changes in expression levels. These results provide implications for the molecular mechanisms behind non-uniform phenotypic variability. The observed transcriptional bias was consistent across diverse conditions, including adaptive evolution, and was closely related to crucial functions for coping with fluctuating environments, suggesting its pervasive impact on both genetic and nongenetic adaptation to environments.

A major limitation of our study is that it focused solely on the transcriptome regulatory networks of a single strain of *E. coli* (MG1655 including its closely-related strains). Currently, most of the reliable knowledge about transcriptional regulatory networks in *E. coli* is heavily restricted to this model strain. Ideally, gathering comprehensive information on the networks of different strains of *E. coli* or other bacterial species with different configurations of transcriptional regulatory networks would enable us to explore how transcriptional variability depends on their transcriptional regulatory networks. This future exploration would also require many transcriptomic profiles under different environmental conditions for each strain to capture the transcriptional variability unique to each. This extension would further underscore the impact of transcriptional regulatory networks on the magnitude and directionality of transcriptional variability.

## Methods

### Bacterial strains and media

We constructed a mutator strain of *E. coli* MG1655 by deleting the *dnaQ* gene. Deletion was performed using conventional λ RED recombination^49^. In brief, a kanamycin resistance gene flanked by FLP recognition sites was amplified from *E. coli* BW25113 *ΔmoaA::kan*, a strain of the Keio collection^50^, by PCR with chimeric primers containing homology arms for the *dnaQ* region. We used the same chimeric primers as those used to knock out *dnaQ* in a previous study^50^. The PCR fragment was used for lambda RED-mediated homologous recombination with pKD46. The obtained strain after obviating pKD46 was called as MG1655 *ΔdnaQ* and used for the MA experiment. We used YENB broth^51^ supplemented with 60 mM arabinose (FUJIFILM Wako, Osaka, Japan, 012-04581) and 50 μg/mL ampicillin (FUJIFILM Wako, Osaka, Japan, 012-23303) to prepare electrocompetent cells for 11 RED recombination. LB agar plates (Miller, BD Difco, 244620) supplemented with 25 μg/mL kanamycin (FUJIFILM Wako, Osaka, Japan, 115-00342) were used as the selective media after electroporation. We used LB agar and broth for the MA experiments and transcriptome analysis of the mutants, respectively.

### MA experiments

The MA experiments were performed as previous studies^30^. In brief, we used two independent colonies of MG1655 *ΔdnaQ* as founder strains, founder A and B, to alleviate a potential bias of the mutations before the MA procedure that might affect the accumulated mutations of the descendants during the MA procedures^52^. We initially established six independent lineages for each founder in preparation for extinction during MA experiments. The MA experiments were conducted by transferring randomly chosen single colonies to fresh LB agar plates daily. The plates were incubated at 32 °C. The MA procedure was repeated for 30 times. The MA lineages after the 10th and final rounds were stored at – 80 °C. Three randomly selected three MA lineages from each founder were subjected to transcriptome analyses (**Supplementary Data 2**).

### Transcriptome profiling using microarray technology

The detailed procedure for transcriptome profiling has been described in previous study^31^. All strains obtained by the MA experiments were grown in 200 μL of LB medium in 96-well microplates at 32 °C. The cell density was monitored using a 1420 ARVO microplate reader (PerkinElmer Inc., Waltham, Massachusetts, USA). The cell broth was sampled at the exponential phase, when OD600 reached in the 0.072–0.135 range. The cell broth was immediately added to an equal volume of ice-cold ethanol containing 10% (w/v) phenol. The cell pellets were obtained by centrifugation at 20,000 × g at 4 °C for 5 min. Total RNA was extracted using an RNeasy Mini kit with on-column DNase I digestion for 15 min (Qiagen, Hilden, Germany) according to the manufacturer’s protocol. Purified RNA samples were analyzed for quality control using an Agilent 2100 Bioanalyzer and an RNA 6000 Nano kit (Agilent Technologies). Microarray experiments were performed using a custom-designed Agilent

8 × 60 K array for the *E. coli* W3110 strain, comprising 12 probes for each gene. Purified total RNA samples (100 ng) were labeled with Cyanine3 (Cy3) using a Low Input Quick Amp WT labeling kit (One-color; Agilent Technologies). The Cy3-labeled cRNA samples were checked for amount (>825 ng) and specific activity (>15 pmol/μg) using a NanoDrop ND-2000 (Thermo Scientific). Subsequently, the labeled cRNA (600 ng) was fragmented and hybridized to a microarray for 17 h while rotating at 10 rpm at 65 °C in a hybridization oven (Agilent Technologies). The hybridized microarray was washed and scanned according to the manufacturer’s instructions. Microarray images were analyzed using Feature Extraction version 10.7.3.1 (Agilent Technologies). Duplicate biological data of the founder strains under the same culture conditions were obtained to confirm technical noise. The compendium of transcriptome data obtained from the MA experiment is called the Mut dataset.

### Compilation of the TRN in *E. coli*

We compiled the reported TRN using all interactions with experimental evidence from RegulonDB 11.0^38^ and EcoCyc^39^ for both transcription factors and sigma factors (**Supplementary Data 7**). For transcription factors, the mode of regulation (activation or repression) was also included. We set the mode of regulation by the sigma factor to activation. When multiple modes were reported for a given regulatory mechanism between a transcription factor and its target gene, they were designated as unknown.

### Analysis of network architecture of TRN using bow-tie decomposition

First, we analyzed the architecture of TRN in *E. coli* based on bow-tie decomposition^42^. We followed the formal definition of the bow-tie structures proposed by Yang *et al*^41^. Their bow-tie structure has seven sectors. The three main sectors are IN, OUT, and SCC. The SCC sector consists of the largest subnetwork in which each node (gene or transcriptional regulator) is reachable by any other node, following the direction of regulation (activation and/or inhibition). The IN sector represents a group of nodes that are not included in the SCC but can reach nodes in the SCC. The OUT sector represents a group of nodes not included in the SCC that can be reached by the nodes in the SCC. The remaining nodes, other than those in the three main sectors, can be further categorized into TUBES, INTENDRILS, OUTTENDRILS, and OTHERS, according to reachability.

To understand how the internal and external perturbations of the cells propagate TRN, we embedded links between the internal/external effectors and TRs, as follows. In addition to the bow-tie structure, we also defined the STIMULI block. The STIMULI block does not consist of a gene or regulator, but consists of intracellular and extracellular effectors, such as nutrients, stressors, metabolites, and stimulating transcriptional regulators by allosteric interactions and so forth. We used the compendium of known links between effectors and TRs collected by Martínez-Antonio *et al*^43^. These target regulators are defined as SR. SRs change their regulatory activity depending on whether they bind to effectors. The networks obtained are listed in **Supplementary Data 8**.

### Data analysis of the three main datasets (Env, Evo and Mut)

In addition to the Mut dataset, we analyzed two datasets of transcriptome in *E. coli*. To estimate the variability in the expression levels of *E. coli* MG1655 in response to environmental changes, we used an updated version of PRECISE^25, 26^, which contains the RNA-seq compendium of *E. coli* MG1655 obtained from a single laboratory using a standardized protocol. In particular, the transcriptome profiles of PRECISE 2.0 were downloaded from iModulonDB^53^. We filtered the transcriptome data of any mutant or evolved strains derived from the MG1655 strain or from other strains to avoid mutational effects on transcriptional variance. Consequently, 160 profiles under 76 unique environmental conditions were used (**Supplementary Data 1**). The environmental conditions were mainly combinations of seven base media (LB, M9, W2, CAMHB, Tris-maleic acid minimal medium, M9 without phosphate, RPMI supplemented with 10% LB), 12 carbon sources (acetate, D-ribose, fructose, galactose, glucarate, gluconate, glucose, glycerol, N-acetylglucosamine, pyruvate, sorbitol, xylose), three nitrogen sources (NH4Cl, cytosine, glutamine), two electron donors (O2, KNO3), three trace elements (aebersold trace element mixture, sauer trace element mixture with or without MgSO4), 35 supplements (cytidine, thiamine, leucine, tyrosine, HCl, lactic acid, adenosine, 2, 2′-dipyridyl, FeCl2, NaCl, paraquat, glutathione, adenine, dibucaine, methionine, salicylate, acetoacetate/LiCl, H2O2, LiCl, ceftriaxone, ciprofloxacin, glutamate, glycine, L-tryptophan, meropenem, threonine, trimethoprim-sulfamethoxazole, KCl, phenylalanine, uracil, L-arginine, pyruvate, CuSO4, ethanol, ZnCl2), two temperatures (37 °C, 42 °C), three pHs (5.5, 7, 8.5) and two culture types(batch, chemostat). The filtered compendium was called the Env dataset. To estimate the variability in expression levels during the evolution of *E. coli* in nature, we analyzed the RNA-seq compendium of natural isolates of *E. coli* during growth in the exponential phase in LB broth, where the RNA-seq profiles were obtained in a single laboratory using a common protocol^27^. We analyzed the RNA-seq profiles of 16 natural isolates (MG1655, E1114, HS, H120, E1167, TA447, TA054, E101, TA280, M114, TA249, 41-1Ti9, ROAR8, JJ1886, UT189, S88) that covered all five primary phylogroups of *E. coli* (**Fig. 1b**). The isolates originated from various sources: MG1655 and HS were isolated from *Homo sapiens* feces; TA054, TA447, TA280, M114, TA249, and ROAR8 were isolated from the feces of *Isoodon obesulus*, *Bettongia lesueur*, *Dasyurus viverrinus*, *Burramys parvus*, *Dasyurus hallucatus*, and *Cephalophus sylvicultor*, respectively; H120 and UT189 were isolated from *H. sapiens* urine; 41-1Ti9 was isolated from a *H. sapiens* Crohn’s disease patient; JJ1886 was isolated from *H. sapiens* blood; S88 was isolated from *H. sapiens* cerebrospinal fluid; E1114 and E101 were isolated from water; and E1167 was isolated from soil. The sources of these isolates have been previously detailed.^27^ The compendium of this dataset was called the Evo dataset.

We used 2622 genes that were common to the three datasets (Env, Evo, and Mut) (**Supplementary Data 3**). For all three datasets, the log2-transformed gene expression levels were used in subsequent analyses. Expression levels were subsequently quantile-normalized. The means and standard deviations of the expression levels of each gene were calculated for each dataset. For the Mut dataset, we first calculated the means and variances of expression levels among the MA lineages of the final rounds, which were derived from the same founder. Subsequently, we averaged the mean and variance of the MA lineages derived from different founders. We obtained the means and standard deviations of the expression levels of MA lineages that underwent 30 rounds of the MA procedure. We found that the standard deviations were dependent on the mean expression levels for every dataset, which can confer a confounding effect on the correlation analysis between the datasets. To compensate for this dependency, we calculated the vertical distance of each standard deviation from a smoothed running median of standard deviations, referred to as the DM, following the method outlined in a previous study^54, 55^. For each gene, we calculated the DM for each dataset and named it DMenv, DMevo and DMmut for the Env, Evo, and Mut datasets, respectively. We also calculated variability at the functional category level based on the DM values. Functional categories were defined by Gene Ontologies (GO)^56, 57^. We constructed a subset (82) of GO by removing redundant GO terms based on semantic similarity among GO terms using the GOSemSim^58^, AnnotationDbi^59^, rrvgo^60^ R packages with default parameters. Alternatively, we used the KEGG pathway database^61^. GO terms and KEGG pathways that consisted of fewer than five genes or more than 300 genes were omitted. The mean DM values among the genes belonging to a given GO term or KEGG pathway were used as a measure of variability at the functional category level **(Supplementary Data 5**,**6**).

Gene set enrichment analysis (GSEA)^35^ was performed using the clusterProfiler R package^62^. We used the refined subset of GO terms and KEGG pathways as functional categories. We filtered out the GO terms and KEGG pathways that consisted of fewer than five genes or more than 300 genes. For GSEA of transcriptional regulators, we also filtered out the transcriptional regulators that regulated fewer than five target genes or more than 1000 genes. The q-values were calculated using the Benjamini-Hochberg (BH) procedure^63^. Transcriptional regulators whose target genes were significantly enriched at higher DM in each dataset were called TRenv, TRevo and TRmut for the Env, Evo, and Mut datasets, respectively. The 13 transcriptional regulators present in all three groups were called TRcom.

The edge density of a given gene set belonging to TRN was calculated as the total number of edges (i.e. regulations) within the gene set divided by the total number of possible edges within the gene set. Self-loops (i.e. autoregulations) were ignored. To compare the edge density within TRcoms with that within randomly chosen regulators, we generated 10,000 subsets by drawing the same number of TRcoms (i.e. 13) transcriptional regulators at random from a pool of transcriptional regulators. The transcriptional regulators that regulated fewer than five target genes were omitted.

PCA was conducted using the prcomp function in R. For the Mut dataset, we used all transcriptional profiles, including isolates at rounds 10 and 30, as well as the founders. In PCA, the *k-* th principal component, *PCk*, is represented as a linear combination of loadings *a*, and expression levels *x*, as follows:

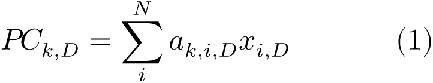

where *D*, *N* and *i* represent the datasets (ex. Env), total number of genes, and *i*-th gene, respectively. Correlation analysis was performed for the loadings of the top three PCs, that is, PC1–3 between the datasets. The correlation was evaluated using absolute values of the Spearman’s rank correlation coefficients (R), as the signs of the PCs are arbitrary.

### Data analysis of the Rwr dataset

The Rwr dataset was derived from transcriptome profiles reported by Baumstark *et al* ^45^. In the Rwr dataset, *E. coli* TOP10 (K12) strains, which harbor synthetic shuffled operons ^44^ (referred to as rewired strains), were cultured under identical environmental conditions. Transcriptome profiles were obtained using the DNA microarray technique (**Supplementary** Fig. 13a,b). A total of 2442 genes, shared across the three main datasets, were extracted. The expression profiles were log2-transformed and quantile-normalized. To facilitate comparison with the Env dataset, which exhibits high reproducibility, transcriptional profiles that showed relatively low quality (less than 0.95 in mean Spearman’s R between biological triplicates) were excluded from the Rwr dataset. The transcriptional profiles were subsequently averaged between the triplicates for each rewired strain. We used a computationally generated random dataset, following a multivariate normal distribution, to confirm that this filtration and averaging process yielded high reliability, comparable to the Env dataset. This filtration resulted in transcriptional profiles of 28 rewired strains (**Supplementary** Fig. 13c).

### Data analysis of the other omics datasets (Strs_hp, Strs_he, Strs, Prtn and Noise_ch)

The Strs_hp and Strs_he datasets were obtained from transcriptome profiles reported by Horinouchi *et al*^64^. In Strs_hp, *E. coli* MDS42^65^, a derivative of MG1655, was cultured under different stress conditions, and transcriptome profiles were obtained using the DNA microarray technique (**Supplementary** Fig. 19a). The transcriptome profiles obtained from the same stressors used in the Env dataset were removed. Transcriptome profiles for eight stressful conditions (cobalt chloride, sodium carbonate, l-malate, methacrylate, crotonate, methylglyoxal, *n*-butanol and cetylpyridinium chloride) and one stress-free condition (modified M9 medium) were analyzed (**Supplementary Data 9**). Thus, the difference in transcriptome profiles within the obtained dataset, called the Strs_hp dataset, represents transcriptional variability in response to environmental perturbations. For Strs_he, transcriptome profiles were obtained for mutants resistant to each of the different stressors cultured under stressor-free conditions using DNA microarrays (**Supplementary** Fig. 19b). These resistant mutants were obtained through adaptive laboratory evolution experiments using MDS42 as the ancestral strain (**Supplementary** Fig. 19c). We removed the transcriptional profiles obtained from the mutants that experienced the same stressors used in the Env dataset. We analyzed the transcriptional profiles of the mutants obtained under the eight stress conditions described above (**Supplementary Data 9**). Thus, the difference in transcriptional profiles within the resultant dataset, called the Strs_he dataset, reflected the transcriptional variability against genetic mutations accumulated through adaptive evolution. Both datasets contained expression levels of 3741 genes (**Supplementary** Fig. 19d).

The Strs dataset was obtained from the transcriptome profiles reported by Maeda and Iwasawa *et al*^31^. In the Strs dataset, the different MDS42 mutants isolated from adaptive laboratory evolution experiments in the presence of different stressors were subjected to DNA microarray analysis (**Supplementary** Fig. 22a–c). The evolutionary experiment was aimed at obtaining resistance mutants against each of the different stressors using MDS42 as an ancestral strain. The transcriptome profiles of the mutant and ancestral strains were obtained under stressor-free conditions, similar to those of Strs_hp. We removed the transcriptome profiles of mutants that evolved under the same stressors used in the Env dataset (example. hydrogen peroxide, meropenem). We analyzed the transcriptional profiles of the mutants obtained under the 47 stress conditions as detailed in **Supplementary Data 10**. Thus, the difference in transcriptome profiles within the obtained dataset, called the Strs dataset, contained transcriptional variability against genetic mutations that accumulated through adaptive evolution.

For the Strs_hp, Strs_he and Strs datasets, the mRNA expression levels obtained using DNA microarrays were log2-transformed and quantile-normalized, similar to the Mut dataset.

The Prtn dataset was obtained from proteomic profiles reported by Schmidt *et al*^66^. Proteomic profiles with the growth rate were determined for *E. coli* BW25113 cultured under 22 different environmental conditions (including different carbon sources). The difference in proteomic profiles within the resultant dataset, called the Prtn dataset, represented the proteomic variability (2002 proteins) in response to environmental perturbations (**Supplementary** Fig. 17). The absolute abundance of proteins (in copies/cell) was log2-transformed.

For all four datasets (Strs_hp, Strs_he, Strs, and Prtn), the means and standard deviations of the expression levels of each gene were calculated across either the mutant or environment in each dataset (**Supplementary** Figs. 17,19,**22**). All datasets showed that the standard deviations depended on the mean expression levels. To compensate for this dependency, we calculated the DM values for each gene as the vertical distance of each standard deviation from a smoothed running median of the standard deviations for each dataset as detailed above. Consequently, we obtained DMstrs_hp, DMstrs_he, DMstrs and DMprtn for Strs_hp, Strs_he, Strs, and Prtn, respectively. PCA was performed as described previously (**Supplementary** Figs. 18,21,**23**).

The Noise_ch dataset was obtained from the cell-to-cell heterogeneity of protein expression levels reported by Taniguchi *et al*^67^ (**Supplementary** Fig. 24). Briefly, a library of yellow fluorescent protein (YFP) fusion strains of *E. coli* K12 was cultured under identical conditions, and each chromosomal copy of the gene was labeled with YFP fusion. YFP fluorescence was measured using microscopy at the single-cell level, which yielded the fluorescence distribution for each gene (1018 genes). For each gene, the protein noise was defined as the variance divided by the square of the mean, which was identical to the coefficient of variation (CV^2^)^67^. Therefore, the protein noise in the Noise_chr dataset represents isogenic cell-to-cell heterogeneity in protein expression levels. It is well known that protein noise is mainly composed of two sources: intrinsic noise dominating lowly expressed proteins and extrinsic noises dominating highly expressed proteins. Owing to the Poisson-like statistical nature of intrinsic noise, the protein noise depends on the mean expression levels of the lowly expressed genes (**Supplementary** Fig. 24c). To compensate for this dependency, we calculated the DM values for each gene as the vertical distance of each protein noise (log10-transformed CV^2^) from a smoothed running median of the protein noise; this conventional compensation is widely used for protein noise^54, 68^. Consequently, we obtained DMnoise_ch, as a measure of variability against stochastic perturbations (**Supplementary** Fig. 24d). Using the noise limits defined by Taniguchi *et al*., we determined the expression level at which the intrinsic noise limit was equal to the extrinsic noise limit. Genes with expression levels below this threshold were defined as lowly expressed, whereas those with expression levels beyond this threshold were defined as highly expressed. We compared DMnoise_ch with the DM values of the other datasets for lowly and highly expressed genes separately.

### Data visualization

All figures were generated in R^69^ and the following packages were used. The phylogenetic tree of the 16 natural isolates of *E. coli* was visualized using the ape^70^ and ggtree^71^ packages. Pairwise phylogenetic distances were based on core genome alignments and were obtained from a previous study^27^. Network visualization was performed using the ggraph^72^, igraph^73^ and tidygraph^74^ packages. The Venn diagram was visualized using the ggVenndiagram^75^, gplots^76^ and venneuler^77^ packages. Other illustrative plots were generated using the ggplot2^78^ and ggpubr^79^ packages.

### Statistics and reproducibility

Two-sided Spearman’s rank correlation coefficients were calculated using the cor.test function in R, specifying the method as Spearman. GSEA was performed with the GSEA function in the clusterProfiler R package. The reproducibility of transcriptome profiles in the Env and Mut datasets was confirmed throught biological replicates (**Supplementary** Fig. 1). No statistical method was used to predetermine sample size.

## Data availability

All microarray data generated in this study have been deposited in the National Center for Biotechnology Information’s Gene Expression Omnibus functional genomics data repository (GEO) under accession number GSE260863 [https://www.ncbi.nlm.nih.gov/geo/query/acc.cgi?acc=GSE260863]. Previously published transcriptome profiles of the Env dataset are available in iModulonDB under accession code PRECISE2.0 [https://imodulondb.org/dataset.html?organism=e_coli&dataset=precise2]. Previously published raw reads in RNA-seq experiments of the Evo dataset are available in ArrayExpress with accession number E-MTAB-9036 [http://www.ebi.ac.uk/arrayexpress/experiments/E-MTAB-9036/], with processed transcriptome profiles available in Rousset *et al.* ^27^ Previously published raw and processed transcriptome profiles of the Strs_hp, Strs_he datasets are available in GEO under accession number GSE89746 [https://www.ncbi.nlm.nih.gov/geo/query/acc.cgi?acc=GSE89746]. Previously published raw and processed transcriptome profiles of the Strs dataset are available in GEO under accession number GSE137348 [https://www.ncbi.nlm.nih.gov/geo/query/acc.cgi?acc=GSE137348]. Mass spectrometry raw data of the Prtn are available in ProteomeXchange Consortium [http://proteomecentral.proteomexchange.org/] under accession number PXD000498 [https://proteomecentral.proteomexchange.org/cgi/GetDataset?ID=PXD000498], with processed proteome profiles available in Schmidt *et al.* ^66^ Previously published protein noise data of the Noise_ch dataset are available in Taniguchi et al. ^67^ Previously published raw reads in scRNA-seq experiments analyzing cell-cycle-dependent changes in gene expression are available in GEO under accession number GSE217715 [https://www.ncbi.nlm.nih.gov/geo/query/acc.cgi?acc=GSE217715], with processed data available in Pountain *et al.* ^33^

## Code availability

Custom scripts used in the manuscript are available at https://github.com/tsuruubi/Variability_E_coli^80^.

## Acknowledgements

The authors thank H. Koike for providing technical assistance with the MA experiments. The authors acknowledge H. Kotani for providing technical assistance with the DNA microarray experiments. This work was supported by, the Japan Society for the Promotion of Science (JSPS) KAKENHI (18H02427, 24K21985 to S.T.;17H06389 to C.F. and S.T.; 22K21344 to C.F.) and the Japan Science and Technology Agency (JST) ERATO (JPMJER1902 to S.T. and C.F.).

## Author Contributions Statement

S.T.: conceptualization, methodology, investigation, visualization, data curation, formal analysis, validation, writing the original manuscript draft, writing—review & editing, fund acquisition., C.F.: resources, writing—review & editing, fund acquisition.

