## Supplementary Fig for "Genetic properties underlying transcriptional variability across different perturbations"

### Supplemental Information for
### Genetic properties underlying transcriptional variability across different perturbations

Saburo Tsuru[1,*] & Chikara Furusawa[1,2,3,*]

[1]Universal Biology Institute, Graduate School of Science, The University of Tokyo, 7-3-1 Hongo, Bunkyo-ku, Tokyo 113-0033, Japan
[2]Department of Physics, Graduate School of Science, The University of Tokyo, 7-3-1 Hongo, Bunkyo-ku, Tokyo 113-0033, Japan
[3]Center for Biosystems Dynamics Research (BDR), RIKEN, 6-2-3 Furuedai, Suita, Osaka 565-0874, Japan

*Corresponding authors:
Saburo Tsuru

Chikara Furusawa


Supplementary Figures
Supplementary Note 1
Supplementary reference

#### Supplementary figures

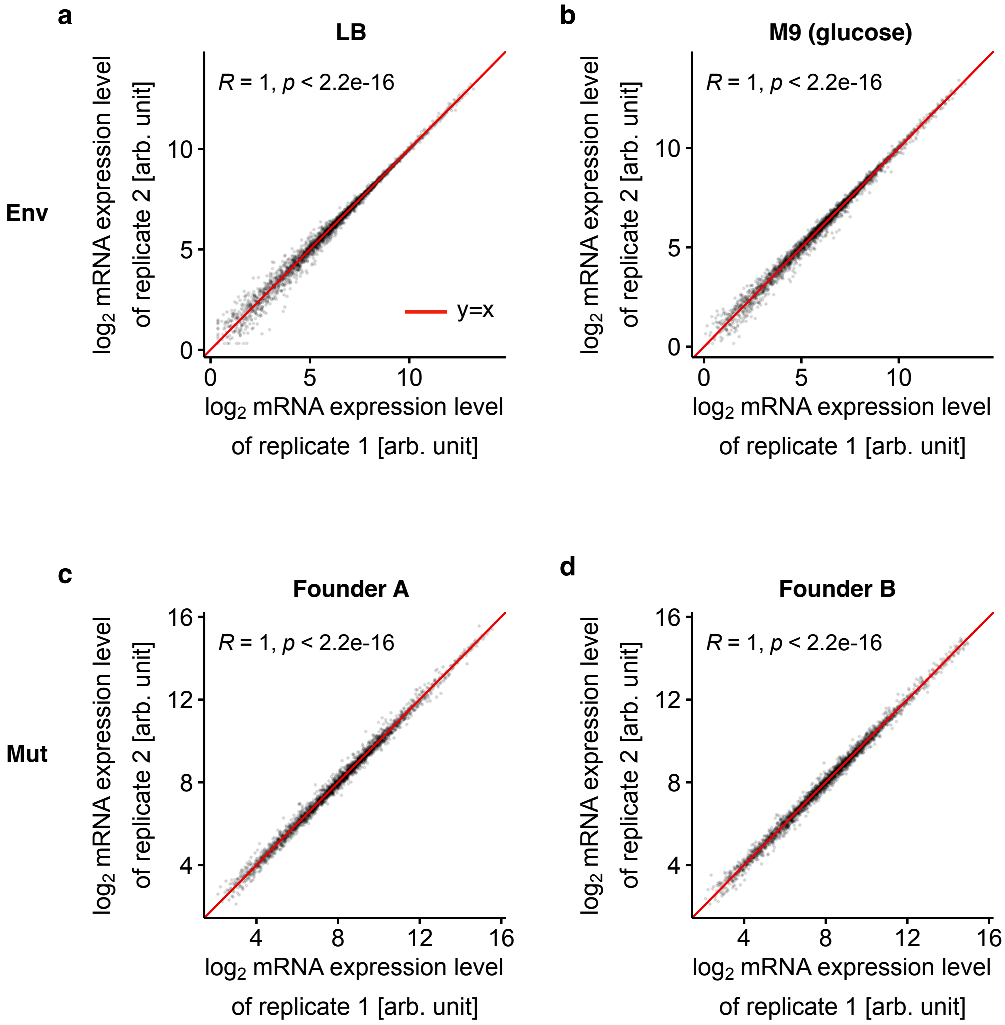


**Supplementary Fig. 1: Reproducibility of transcriptome analysis in Env and Mut.**

**a**,**b**, $Log_2$-transformed and quantile-normalized mRNA expression levels of 2622 genes are shown for two biological replicates (replicates 1 and 2) cultured under a complex rich medium (LB, **a**) and a synthetic minimal medium (M9, **b**) in the Env dataset. **c**,**d**, $Log_2$-transformed and quantile-normalized mRNA expression levels of 2622 genes are shown for two biological replicates (replicates 1 and 2) of founders A (**a**) and B (**b**) in the MA experiment. Each dot represents a different gene. The insets represent Spearman's R and p-values (two-sided). The solid lines represent y= x.

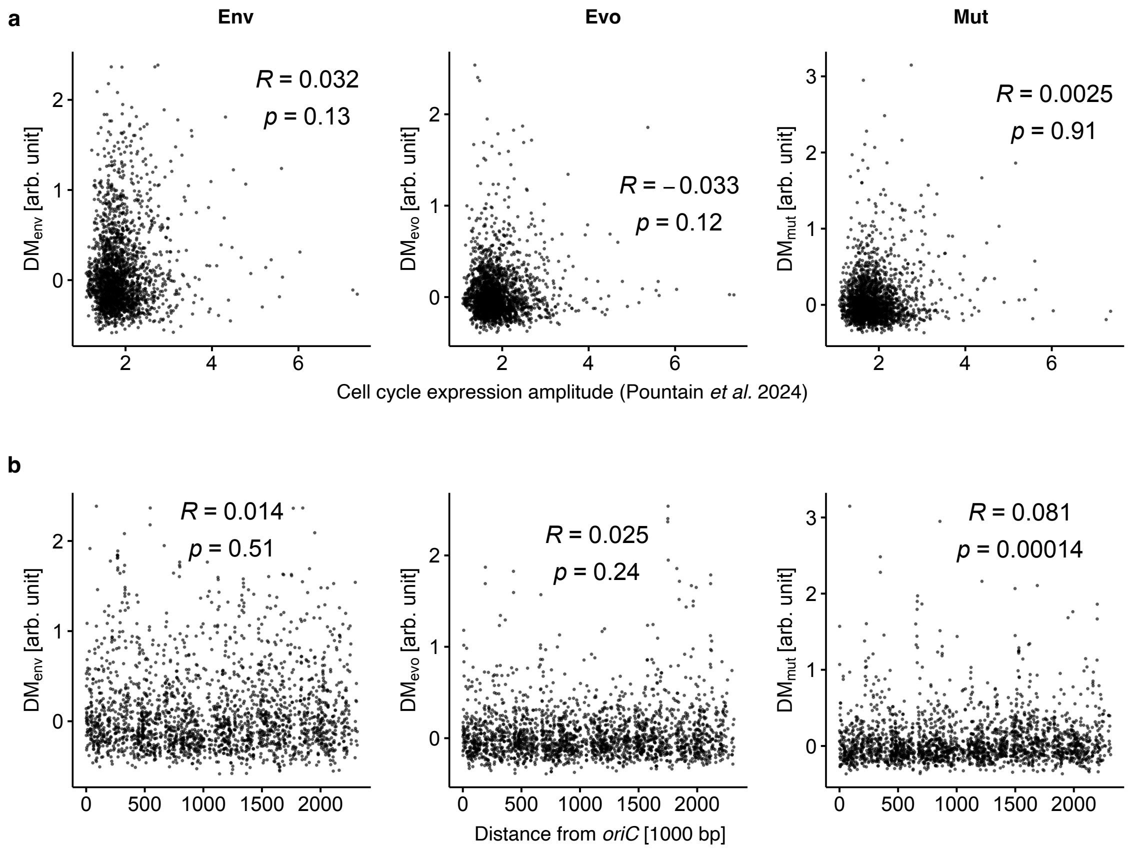


**Supplementary Fig. 2: Relationship between DM values and cell-cycle-associated transcriptional changes.**

**a**, The relationship between DM values and cell cycle expression amplitude. The cell cycle expression amplitude was obtained from Pountain *et al* [1] and is defined as the fold change from trough expression to peak expression in the temporal dynamics of mRNA expression levels within the cell cycle. **b**, The relationship between DM values and the shortest distance from *oriC* on the chromosome of *E. coli* K12 MG1655. The insets represent Spearman's R and p-values (two-sided).

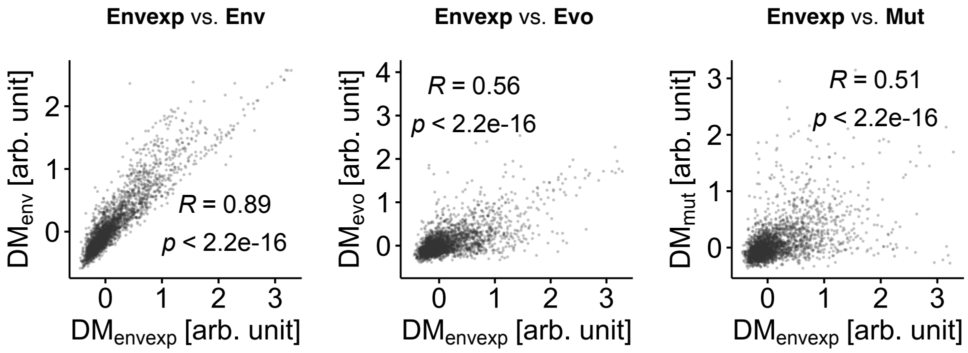


**Supplementary Fig. 3: Relationship between DM values based on exponential growth phase samples.**

Transcriptome profiles obtained during the exponential growth phase were extracted from the original Env dataset to construct the Envexp dataset based on information from the original papers in the PRECISE database. The DM values of genes were calculated from this filtered dataset (78 profiles, 37 unique environmental conditions) and termed $DM_{envexp}$. The relationship between $DM_{env}$, $DM_{evo}$, $DM_{mut}$ and $DM_{envexp}$ is shown. The insets represent Spearman's R and p-values (two-sided). $DM_{evo}$ and $DM_{mut}$ were originally derived from exponential growth phase samples.

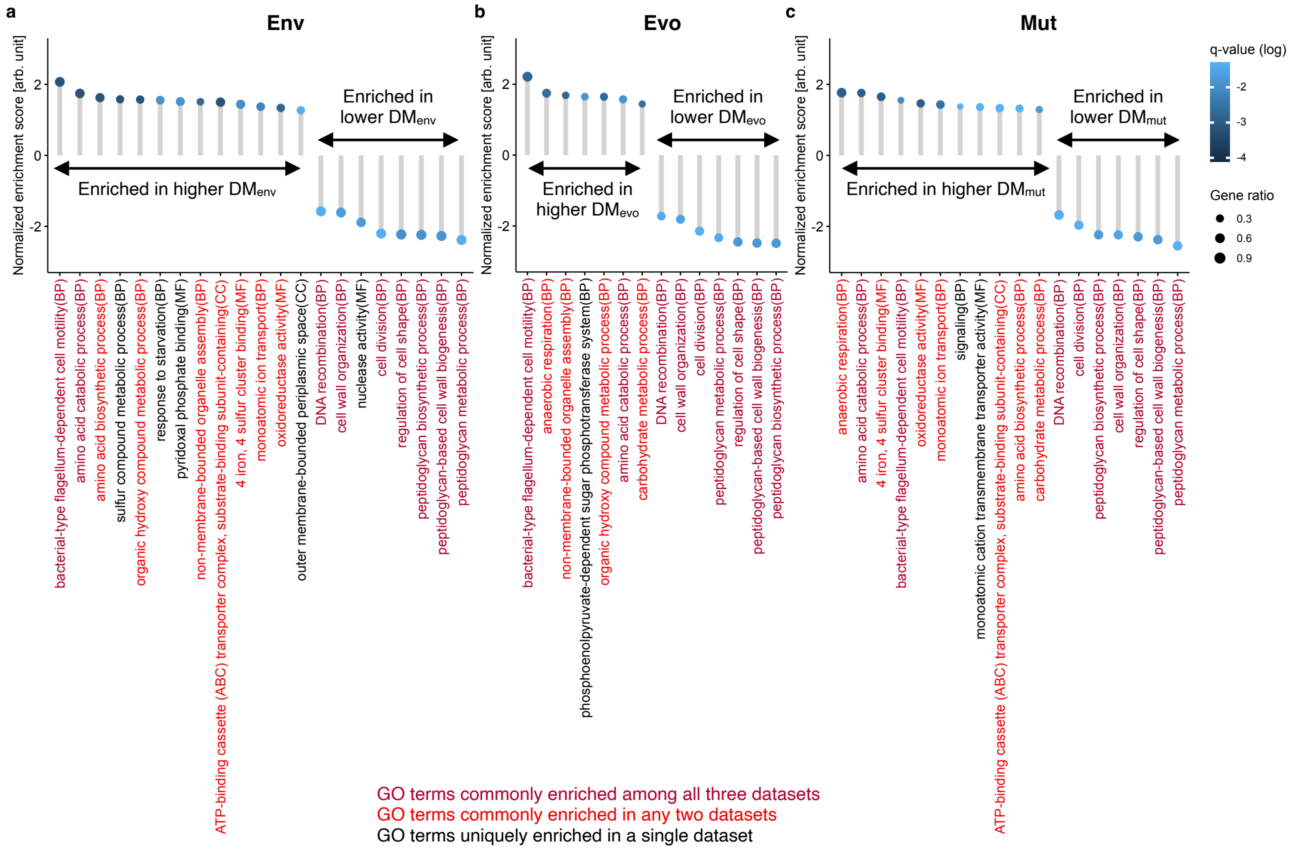


**Supplementary Fig. 4: GO terms enriched in genes with higher and lower DM values.** Normalized enrichment scores of enriched GO terms are shown for Env (**a**), Evo (**b**) and Mut (**c**). The color bar represents the q-values. Only statistically significant GO terms ($q<0.05$) are shown. Positive and negative enrichment scores indicated enrichment at higher and lower DM values, respectively. The circle sizes depict the gene ratio, calculated as the number of core genes contributing highly to the enrichment scores divided by the total number of genes in the respective GO terms. The three domains of GO, namely, biological process (BP), cellular component (CC), and molecular function (MF), are represented. Dark and light red GO terms indicate commonly enriched terms among all three datasets and in two datasets, respectively. The q-values were calculated using the Benjamini-Hochberg (BH) procedure.

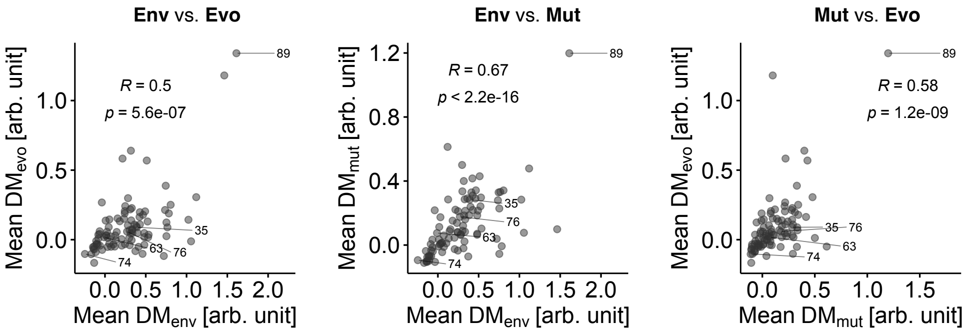


| Label | KEGG ID | Pathway name |
|---|---|---|
| 35 | eco00260 | Glycine, serine and threonine metabolism |
| 63 | eco00790 | Folate biosynthesis |
| 74 | eco00970 | Aminoacyl-tRNA biosynthesis |
| 76 | eco04122 | Sulfur relay system |
| 89 | eco02030 | Bacterial chemotaxis |

**Supplementary Fig. 5: Correlations of DM values at the KEGG functional category levels between dataset pairs.** Each dot represents a KEGG pathway. Only KEGG pathways containing more than four genes and less than 301 genes are displayed. Mean DM was computed by averaging the DM of genes within each KEGG pathway. Representative KEGG pathways are denoted by numbers as specified at the bottom. Spearman's R and p-values (two-sided) are provided within the panels.

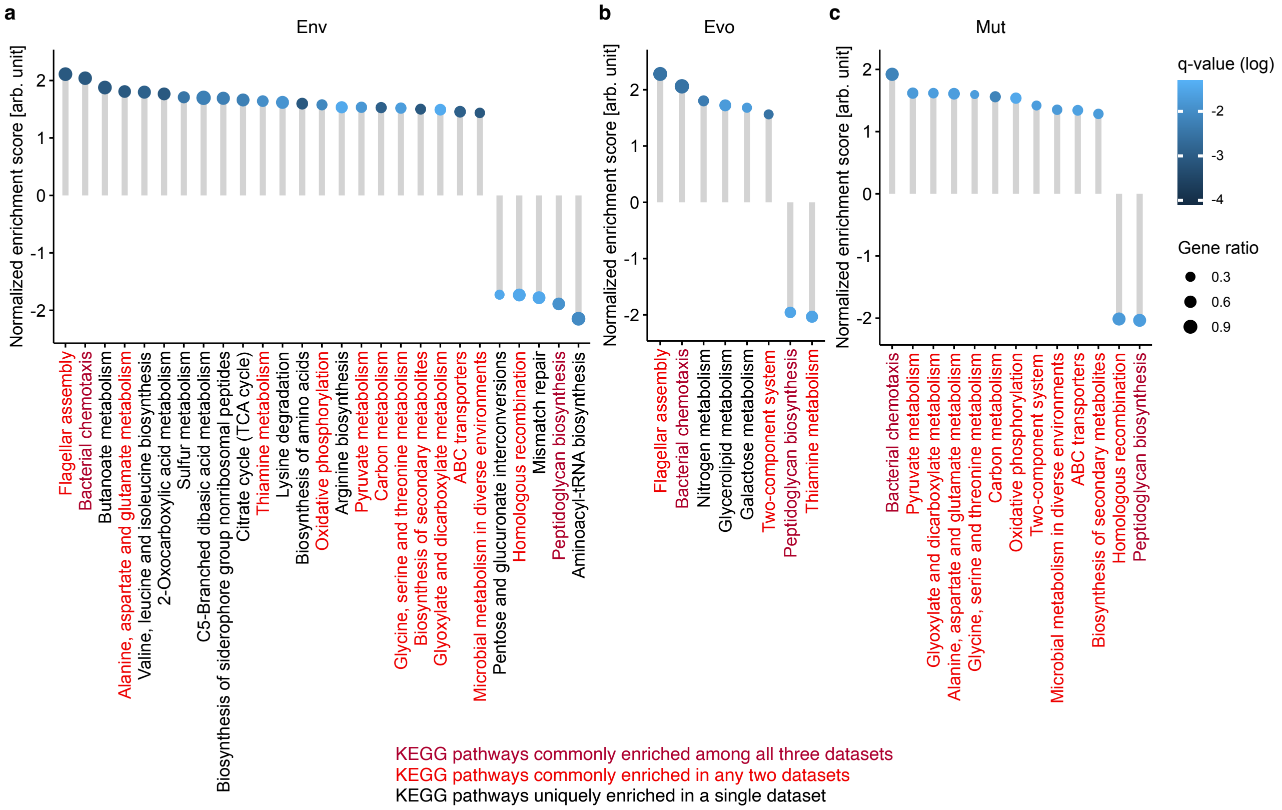


**Supplementary Fig. 6: KEGG pathways enriched in genes with higher and lower DM values.** Normalized enrichment scores for enriched KEGG pathways are presented for Env (**a**), Evo (**b**) and Mut (**c**). The color bar represents the q-values, with only the KEGG pathways displaying statistical significance ($q<0.05$). Positive and negative enrichment scores denote enrichment at higher and lower DM values, respectively. The circle sizes represent the gene ratio, calculated as the number of core genes contributing greatly to the enrichment scores divided by the total number of genes in the focal KEGG pathway. The KEGG pathways shaded in dark and light red represent those commonly enriched across all three datasets or only two datasets, respectively. The q-values were calculated using the BH procedure.

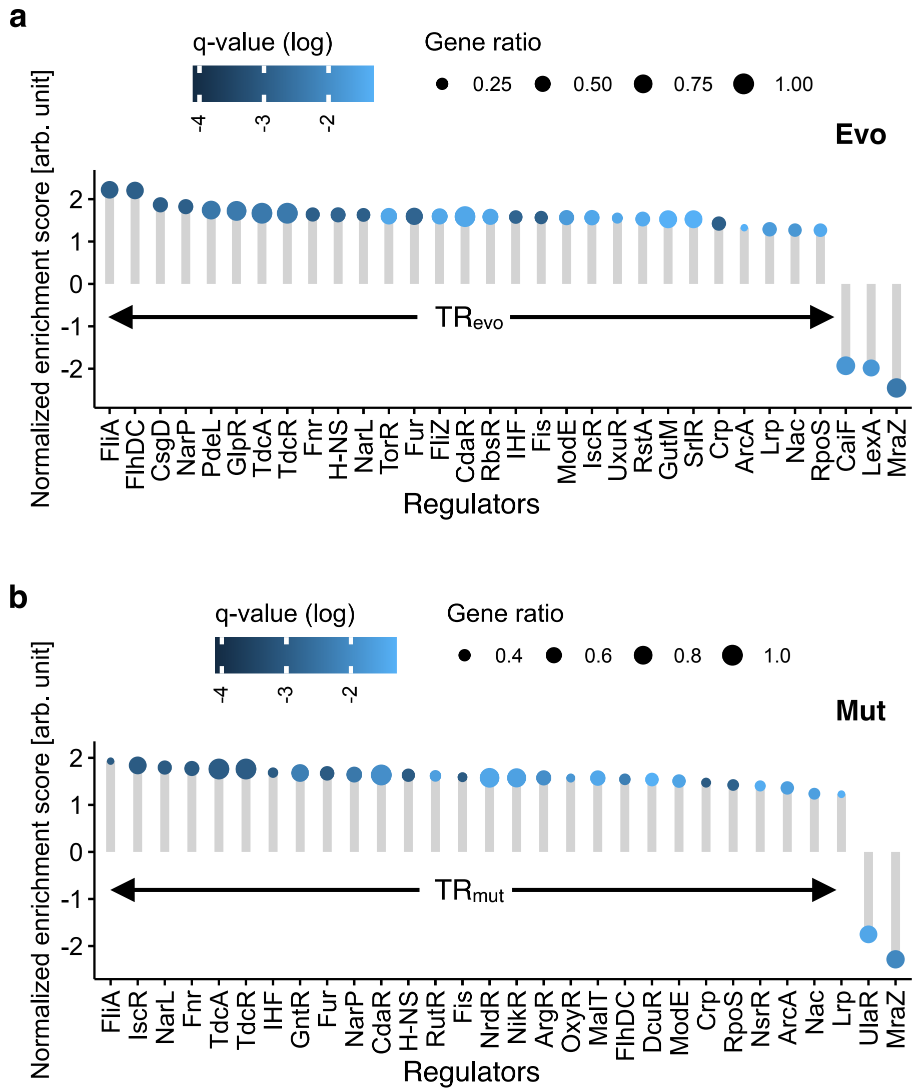


**Supplementary Fig. 7: Transcriptional regulators enriched in the target genes with high and low DM values.**

Normalized enrichment scores for the enriched transcription factors are presented for $DM_{evo}$ (**a**) and $DM_{mut}$ (**b**). Color bars represent the q-values, with only statistically significant regulators shown ($P<0.05$). Positive and negative enrichment scores denote enrichment at higher and lower DM values, respectively. The circle sizes represent the gene ratio, computed as the number of core target genes highly contributing to the enrichment scores divided by the total number of target genes for the focal regulators. Enriched regulators with positive enrichment scores are designated $TR_{evo}$ for Evo (**a**) and $TR_{mut}$ for Mut (**b**). The q-values were calculated using the BH procedure.

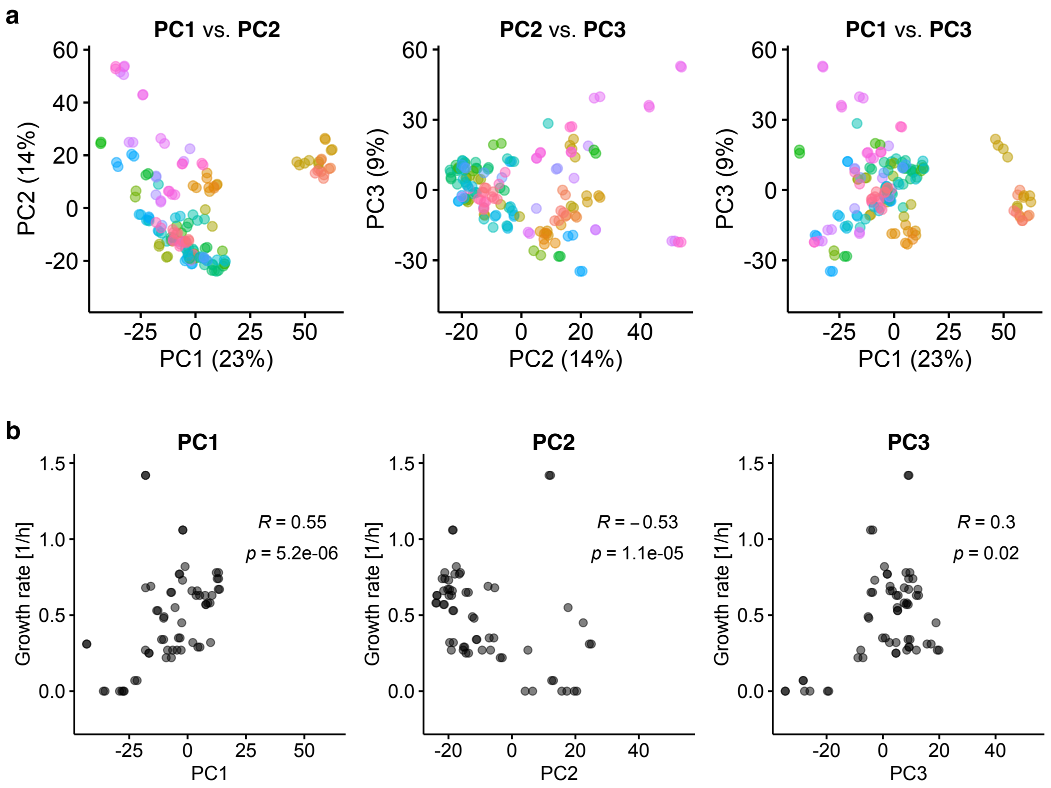


**Supplementary Fig. 8: Top three principal component scores for the Env dataset and the relationship with the growth rate.**

**a**, Pairwise score plots of PC1–3 are shown. Points are colored by environmental condition in Env. **b**, Relationship between PC1–3 and the growth rate under each environmental condition. Spearman's R and p-values (two-sided) are shown in the panels.

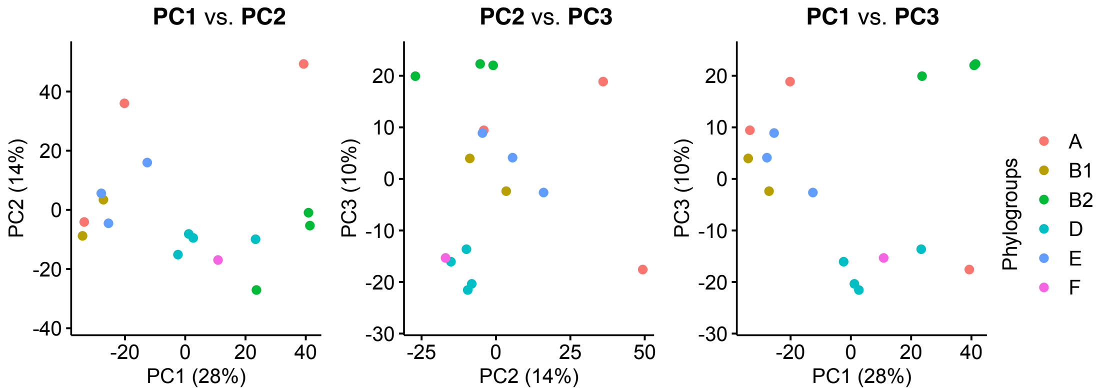


**Supplementary Fig. 9: Top three principal component scores for the Evo dataset.** Pairwise score plots of PC1–3 are shown. Points are colored according to the major phylogroups in *E. coli*.

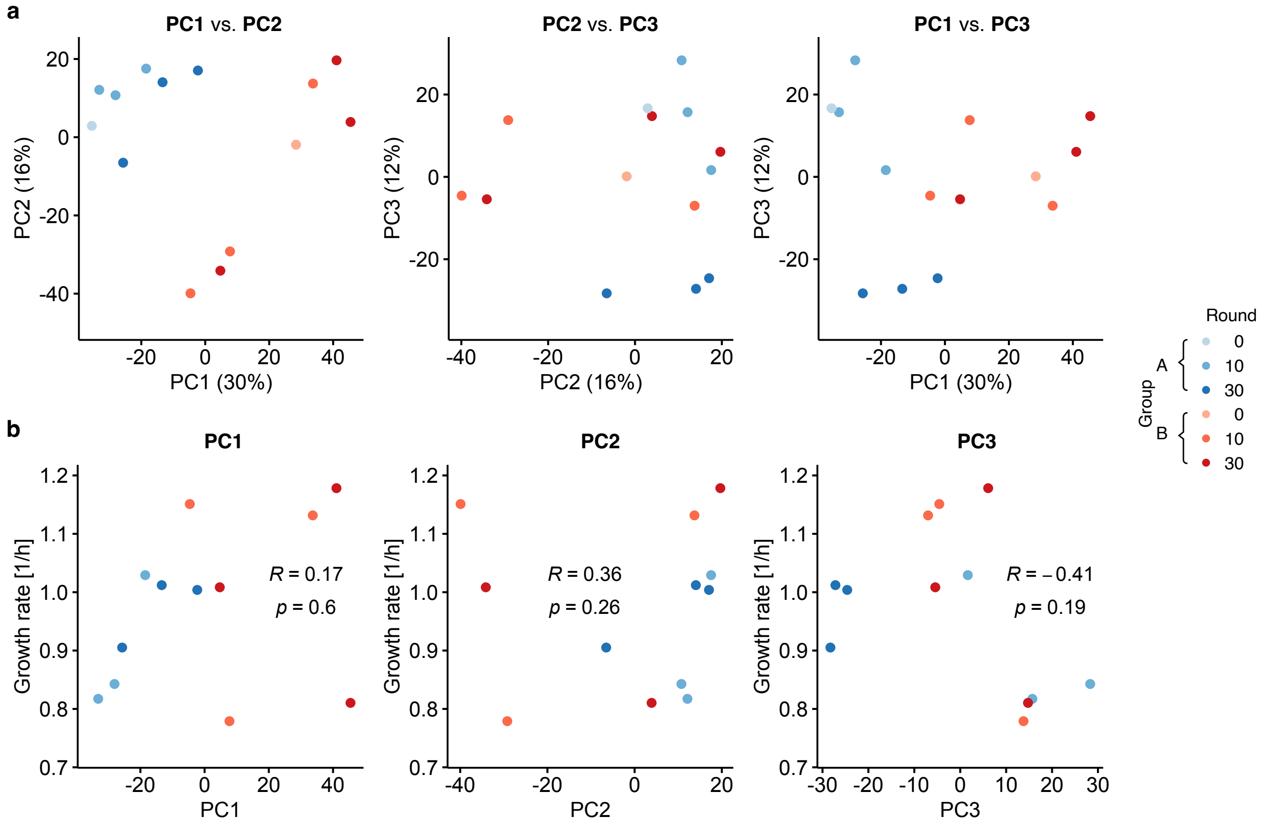


**Supplementary Fig. 10: Top three principal component scores for the Mut dataset and the relationship with the growth rate.**

**a**, Pairwise score plots of PC1–3 are shown. **b**, Relationship between PC1 and 3 and growth rate in LB broth. Points are colored according to the MA group and the round of the MA experiment. Spearman's R and p-values (two-sided) are shown in the panels.

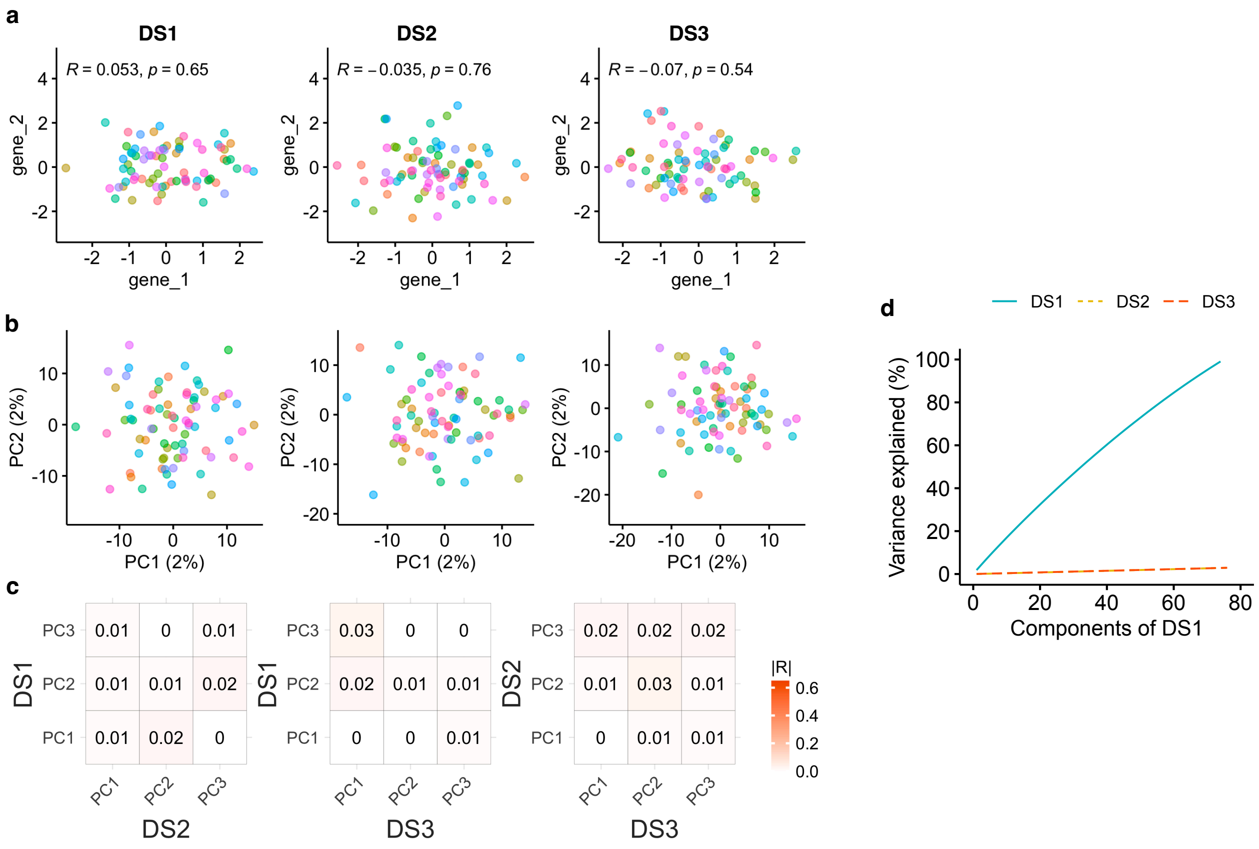


**Supplementary Fig. 11: PCA of the reference datasets comprising computationally generated expression levels.**

Three datasets, termed DS1–3, were computationally generated to serve as a reference group. Each dataset consists of 2,622 genes and 76 conditions, matching the size of the Env dataset. Expression levels were randomly generated for each dataset, following a normal distribution with a mean of 0 and a standard deviation of 1. This ensured the independence of expression levels between genes within and between datasets. **a**, There was no correlation in expression levels between two representative genes (gene_1 and gene_2) for each dataset (DS1 to DS3 from left to right), as designed. Spearman's R and p-values (two-sided) are shown in the insets. **b**, Pairwise score plots of PC1 and PC2 are shown for each dataset. PCA was performed independently for each dataset. Each color represents a different condition. **c**, A correlation analysis was conducted for the loadings of the top three PCs among the three datasets. The color scale and numbers in the tile represent the absolute values of Spearman's R. **d**, The cumulative variance explained by the PCs defined by the DS1 dataset. Overall, the DS1 dataset explained only 3% of the variance in the DS2 and DS3 datasets, respectively.

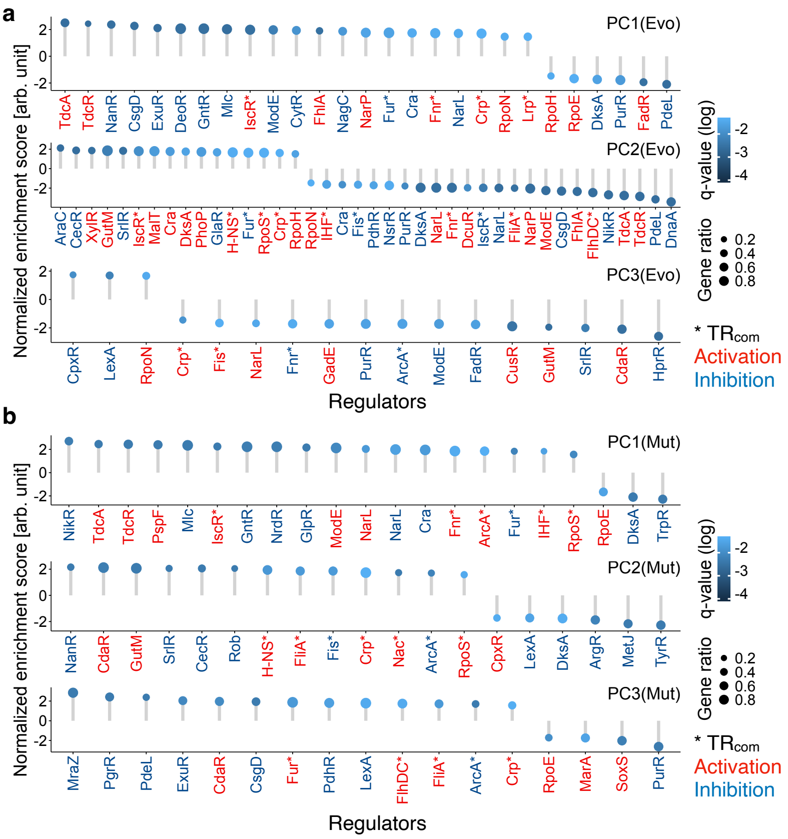


**Supplementary Fig. 12: Transcriptional regulators enriched in the target genes with high contribution to the first three principal components.**

Normalized enrichment scores for the enriched TRs are shown for Evo (**a**) and Mut (**b**). Color bars represent the q-values, with only statistically significant regulators shown (q<0.05). TRs were significantly enriched (q<0.05) in the larger loadings of the first three PCs. GSEA was conducted to identify these regulations and distinguish between activatory (red) and inhibitory (blue) regulations. Positive and negative enrichment scores represented the extent of enrichment in the positive and negative loadings of each PC, respectively. Asterisks represent the regulations by $TR_{com}$.

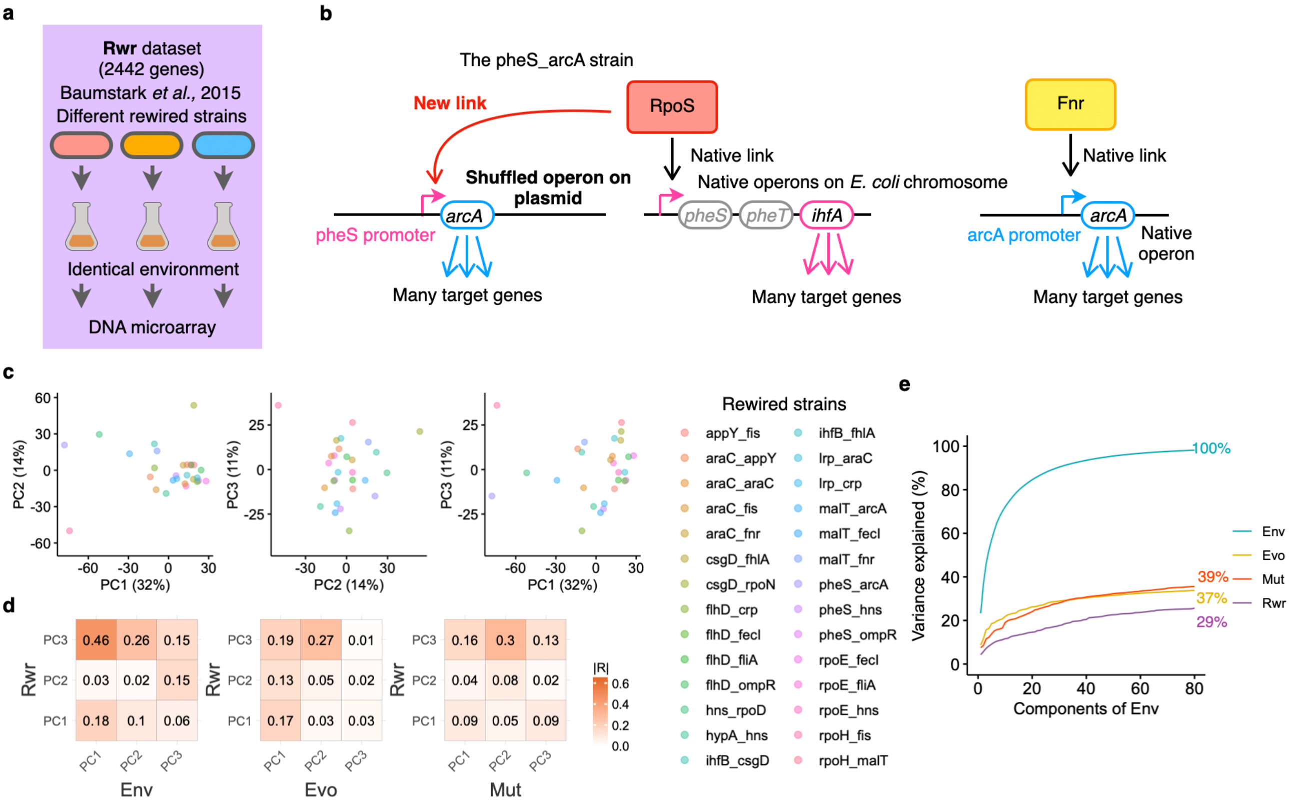


**Supplementary Fig. 13: PCA of the Rwr dataset.**

**a**, A schematic representation of the Rwr dataset. Baumstark *et al*[2] cultured different rewired strains of *E. coli*, originally constructed by Isalan *et al*[3], under identical environmental conditions. Transcriptome profiles were obtained using a DNA microarray. A total of 2,442 genes present in the three main datasets were extracted. Transcriptome profiles were $log_2$-transformed and quantile-normalized. Transcriptome profiles that showed relatively low quality (mean Spearman's R<0.95 between biological triplicates) were removed. The transcriptional profiles were then averaged between the triplicates for each rewired strain. This filtration resulted in transcriptome profiles of 28 rewired strains, termed the Rwr dataset. **b**, A schematic representation of the transcriptional regulation in a representative rewired strain (the pheS_arcA strain). The native promoter region (the pheS promoter) of a global dual transcriptional regulator, *ihfA*, coding for a subunit of IHF, was copied and pasted into a plasmid. The *arcA* gene coding for a global dual regulator ArcA was copied and transcriptionally fused to the plasmid copy of the *pheS* promoter. The native chromosomal copies of these genes and promoters were intact. Consequently, the rewired strain obtained a new regulatory link (curved arrow) between two global regulators, RpoS and ArcA, while the native regulatory links (RpoS to IHF or Fnr to ArcA) were retained. The names of the rewired strains consist of the names of the copied promoters (pheS) followed by the names of the copied ORFs (arcA). **c**, Pairwise score plots of PC1–3 are shown. **d**, A correlation analysis for the loadings of the top three PCs among the datasets. PCA was performed independently for each dataset using the common 2,442 genes shared by the four datasets. The color scale and numbers in the tile represent the absolute values of Spearman's R. **e**, The cumulative variance explained by the PCs defined by the Env dataset. Overall, the Env dataset explained only 29% of the variance in the Rwr dataset (purple).

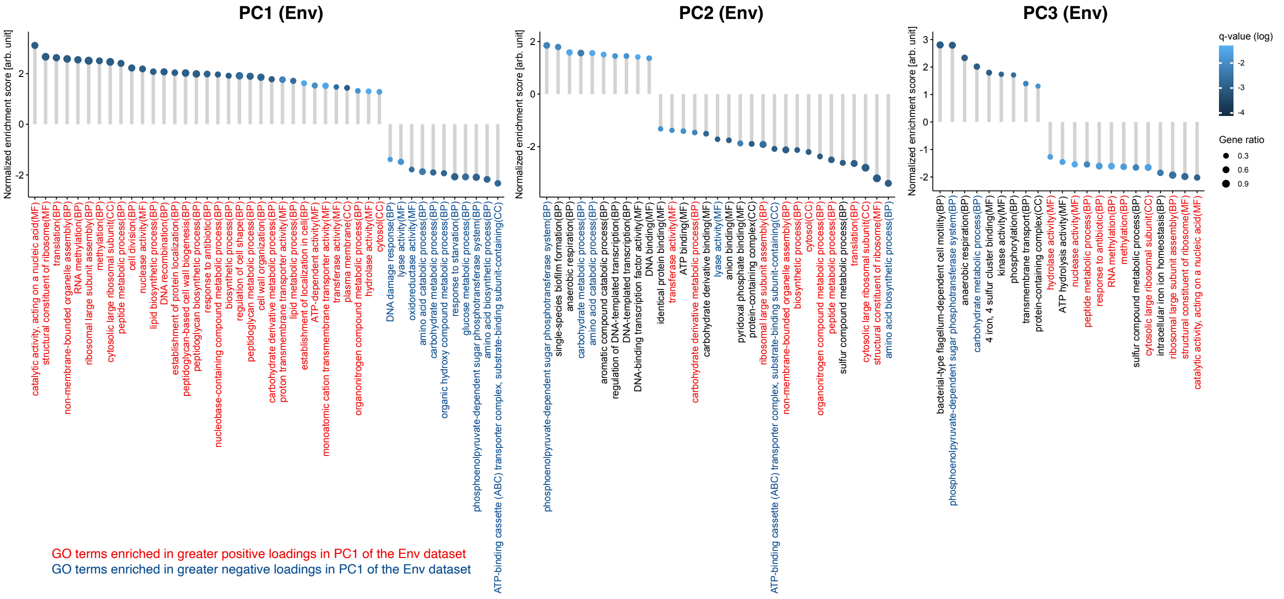


**Supplementary Fig. 14: GO terms enriched in the positively and negatively greater loadings in the first three principal components in Env.**

GSEA was performed to identify the GO terms enriched with greater loadings of PC1–3 in the transcriptome profiles of the Env dataset. The normalized enrichment scores of the enriched GO terms with statistical significance ($q<0.05$) are presented. Positive and negative enrichment scores represented the extent of enrichment in the positive and negative loadings of each PC, respectively. The color bar represents the q-values. The circle sizes represent the gene ratio, calculated as the number of core genes contributing highly to the enrichment scores divided by the total number of genes in the focal GO terms. Biological processes (BP), cellular components (CC), and molecular functions (MF) represented the three GO domains. Red and blue GO terms denote those enriched with greater positive and negative loadings in PC1, respectively.

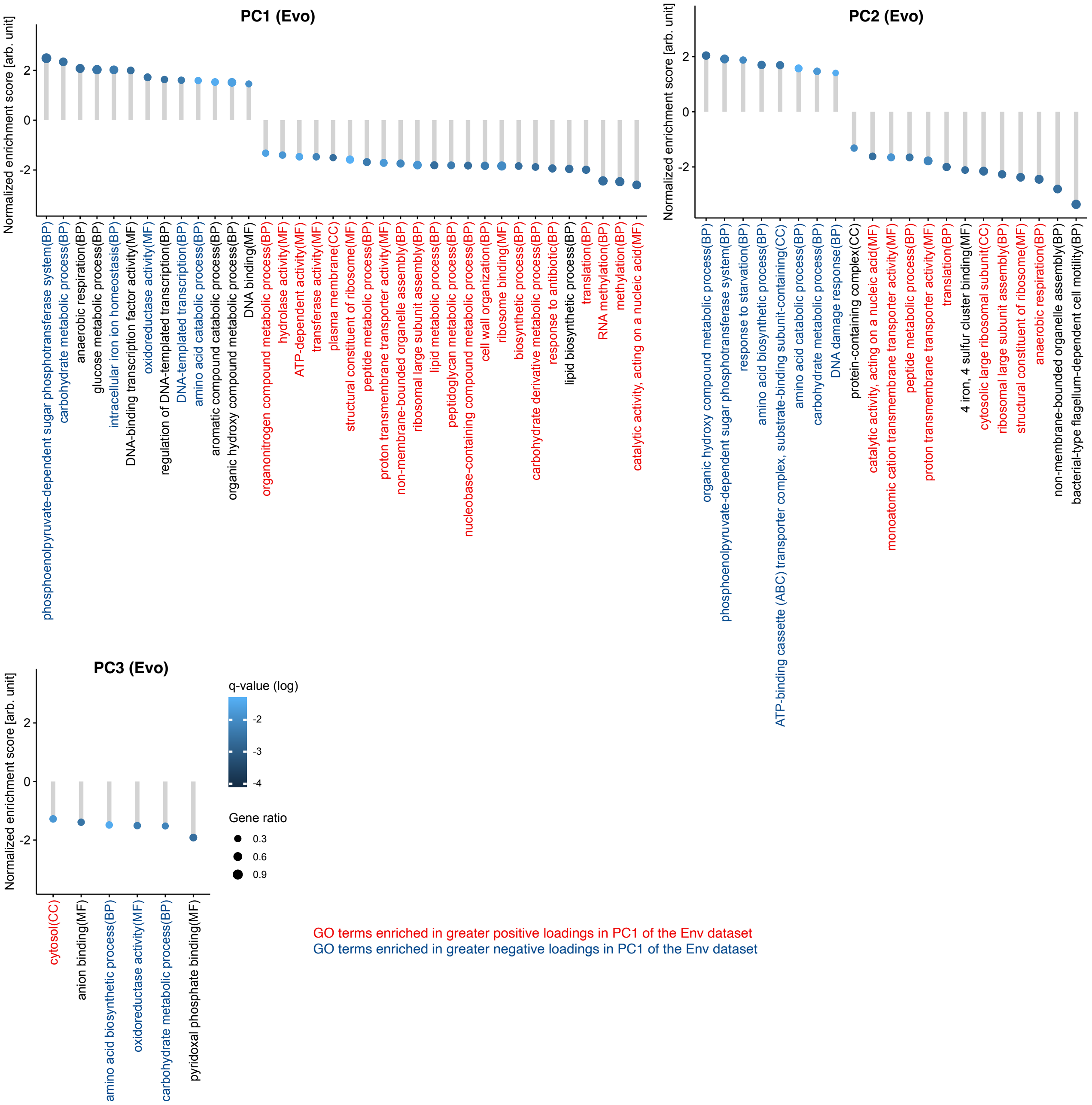


**Supplementary Fig. 15: GO terms enriched in the positively and negatively greater loadings in the first three principal components in Evo.**

GSEA was conducted to identify enriched GO terms with greater loadings in PC1–3 in the transcriptome profiles of the Evo dataset. The normalized enrichment scores of the enriched GO terms with statistical significance ($q<0.05$) are shown. Positive and negative enrichment scores represented the extent of enrichment in the positive and negative loadings of each PC, respectively. The color bar represents the q-values. Circle sizes denote the gene ratio and compute the number of core genes that contribute highly to the enrichment scores divided by the total number of genes in the focal GO terms. Biological processes (BP), cellular components (CC), and molecular functions (MF) represented the three GO domains. Red and blue GO terms represent the GO terms enriched in positive and negative loadings in PC1 of the Env dataset, respectively.

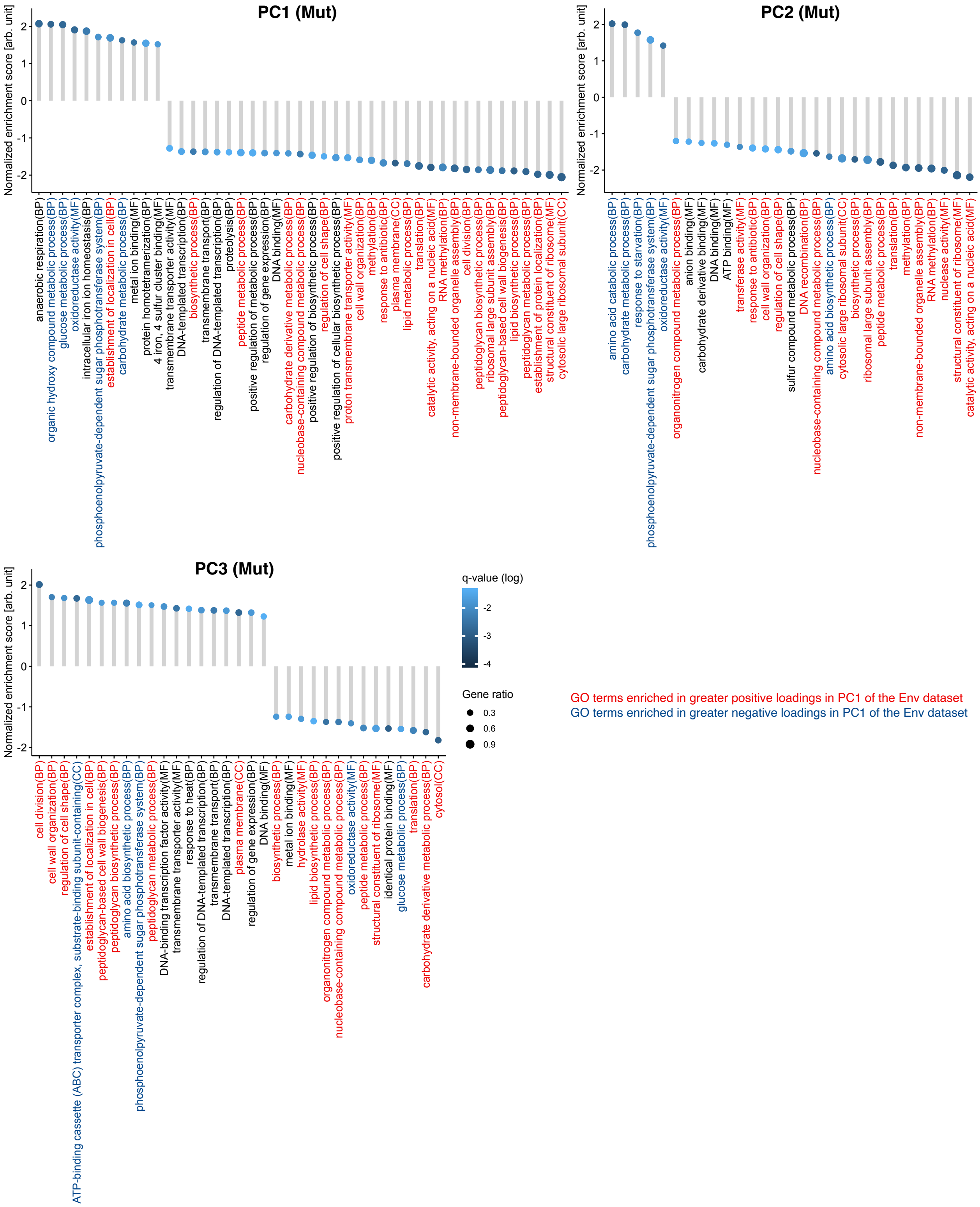


**Supplementary Fig. 16: GO terms enriched in the positively and negatively greater loadings in the first three principal components in Mut.**

GSEA was performed to identify enriched GO terms with greater loadings in PC1–3 in the transcriptome profiles of the Mut dataset. The normalized enrichment scores of the enriched GO terms with statistical significance ($q<0.05$) are presented. Positive and negative enrichment scores represented the extent of enrichment in the positive and negative loadings of each PC, respectively. The color bar denotes the q-values. Circle sizes denote the gene ratio and compute the number of core genes that contribute highly to the enrichment scores divided by the total number of genes in the focal GO terms. Biological processes (BP), cellular components (CC), and molecular functions (MF) represented the three GO domains. Red and blue GO terms represent the GO terms enriched in positive and negative loadings in PC1 of the Env dataset, respectively.

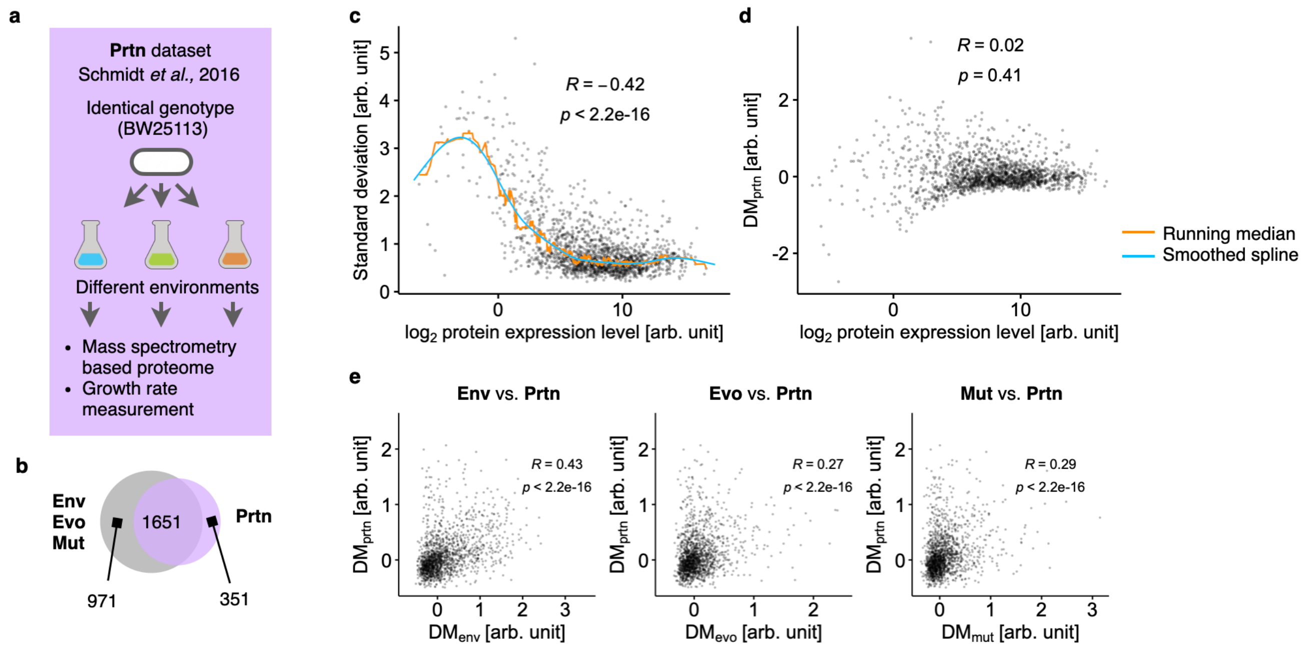


**Supplementary Fig. 17: The DM values in the Prtn dataset.**

**a**, Illustration of the Prtn dataset. Constructed by Schmidt *et al.*[4], the Prtn dataset depicts environmentally induced variations at the protein level. *E. coli* BW25113 was cultured under 22 different environmental conditions, including various carbon sources, and subjected to proteomic analyses. Additionally, growth rates under each condition were measured. **b**, Venn diagram displaying gene overlap between datasets. A total of 1651 genes were common across datasets. **c**, Relationship between mean expression levels and standard deviations. Each grey circle represents a gene. The vertical deviation from a smoothed spline, calculated from the running median of the standard deviations, is denoted as $DM_{prtn}$. **d** Relationship between DM values and mean expression levels. **e**, Correlation of transcriptional variability between the datasets. Pairwise correlations are shown for the DMs between Prtn and the three main datasets (Env, Evo, and Mut) arranged from left to right. Spearman's R and associated p-values (two-sided) are provided in panels **c**–**e**.

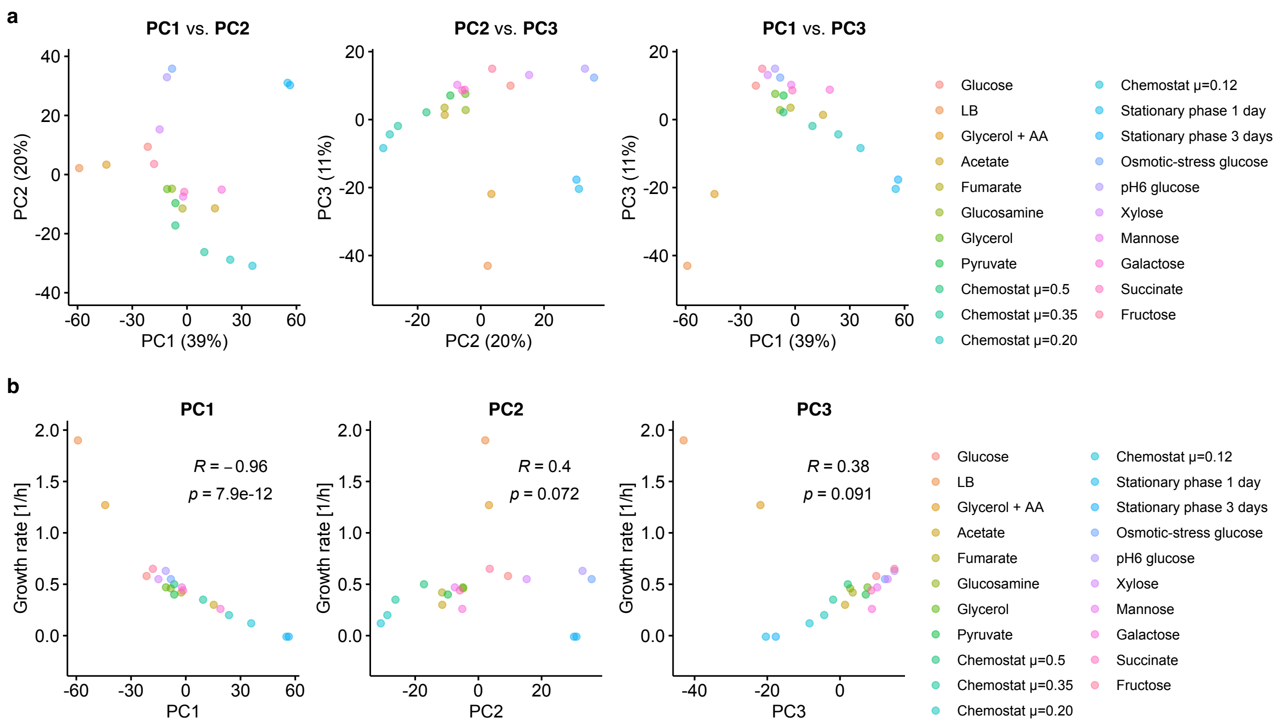


**Supplementary Fig. 18: Principal component analysis of the Prtn dataset.**

**a**, Pairwise score plots of the top three principal components (PCs), PC1–3, for the Prtn dataset. **b**, Relationship between PC1 and 3 and growth rate under each environmental condition. Spearman's R and the associated p-values (two-sided) are provided. The color legend indicates the culture conditions.

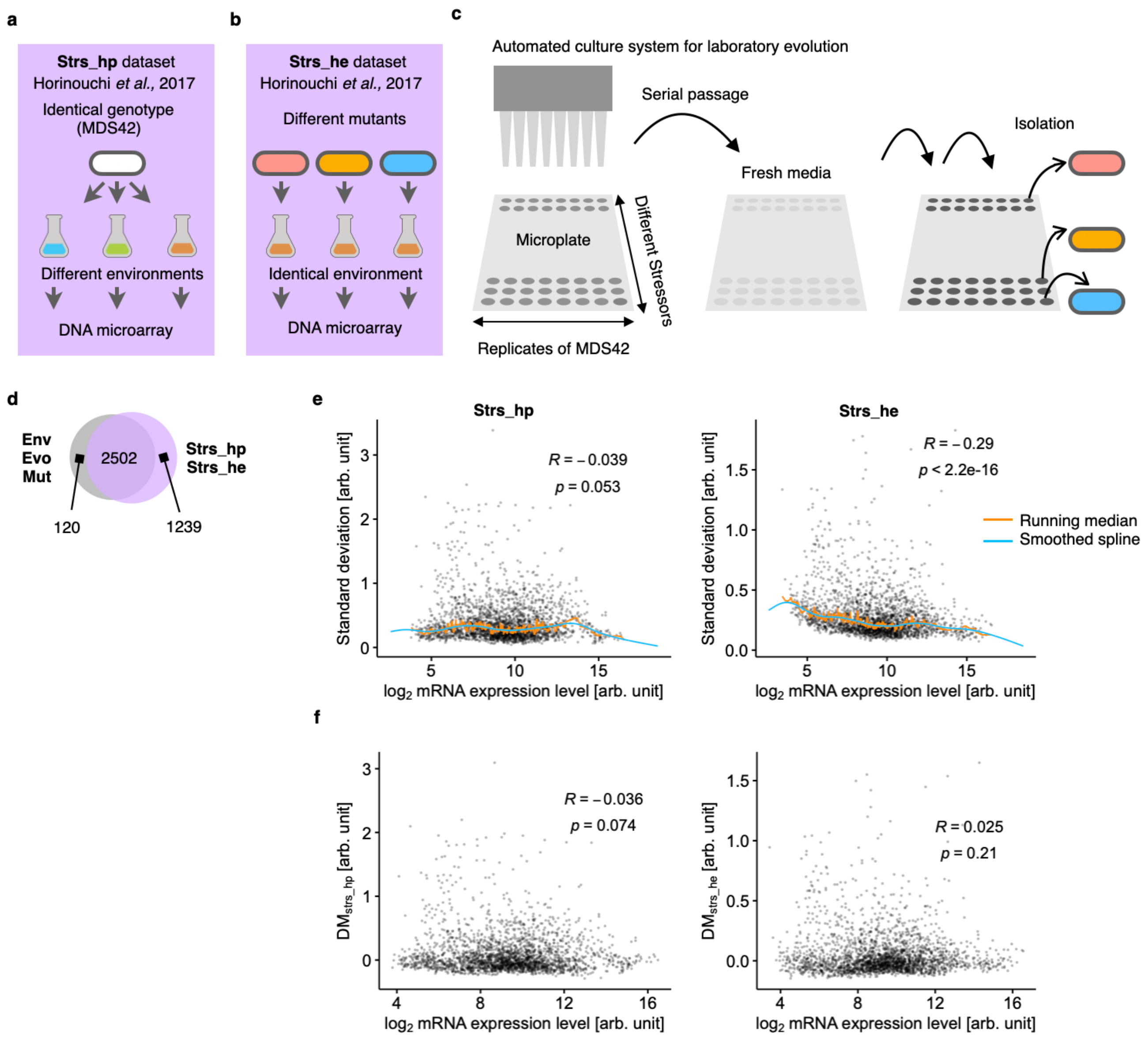


**Supplementary Fig. 19: The DM values in the Strs_hp and Strs_he datasets.**

**a–c** Schematic representation of the experiments on the Strs_hp (**a**) and Strs_he (**b**) datasets. Both experiments were previously conducted in our laboratory[5] using the DNA microarray technique used in this study. **a**, In Strs_hp, *E. coli* MDS42, a derivative of MG1655, was cultured under eight stress conditions (**Supplementary Data 9**), and transcriptome profiles were obtained. We excluded the profiles from the stressors used in the Env dataset, resulting in a dataset that reflected environmentally induced transcriptional variations. **b**, For Strs_he, transcriptome profiles were obtained for the resistant mutants cultured under stressful conditions, representing transcriptional variations induced by genetic perturbations through adaptive evolution. **c**, Evolution experiment to obtain resistance mutants against different stressors using MDS42 as the ancestral strain. Mutants with improved resistance were isolated after serial propagation in stressor-containing medium. Mutants exposed to the stressors used in the Env dataset were excluded. **d**, Venn diagram illustrating shared genes in the datasets. A total of 2502 genes were common. **e**, Relationship between mean expression level and standard deviation. The vertical deviation from a smoothed spline, calculated from the running median of the standard deviations, represents $DM_{strs\_hp}$ for Strs_hp (left) and $DM_{strs\_he}$ for Strs_hp (right). **f**, Relationship between the mean expression levels and DM values. Each grey circle represents a gene. The insets depict Spearman's R and p-values (two-sided).

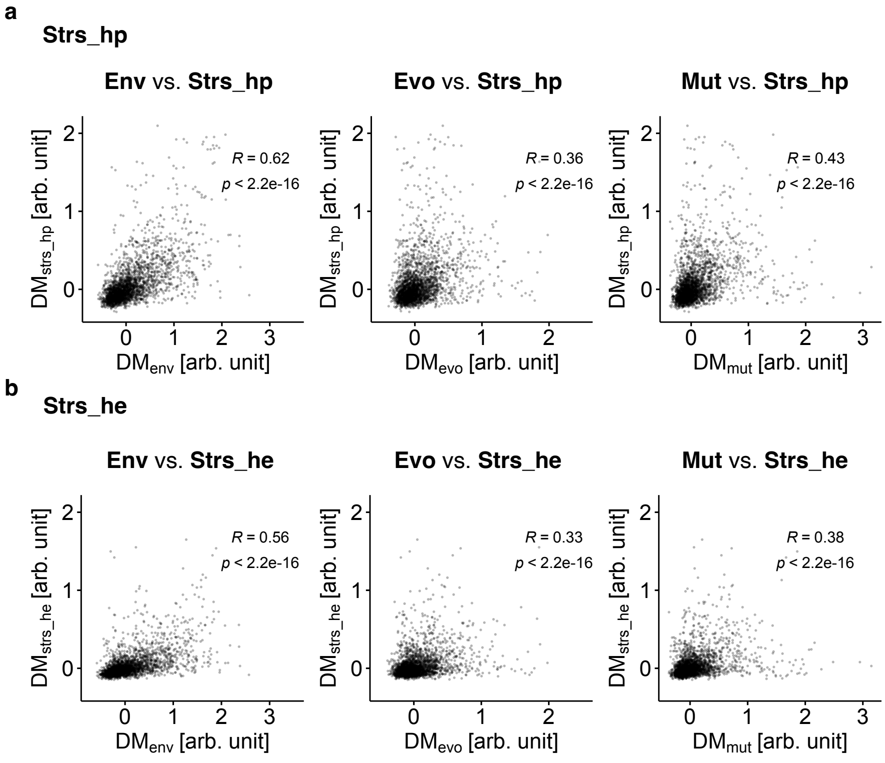


**Supplementary Fig. 20: Positive correlations of transcriptional variability between datasets.** **a**, Pairwise correlations are shown for the DM values between Strs_hp and the three main datasets (Env, Evo, and Mut) arranged from left to right. **b**, Pairwise correlations are shown for the DM values between Strs_he and the three main datasets. In panels **a**,**b**, each dot represents a different gene. Spearman's R and associated p-values (two-sided) are shown.

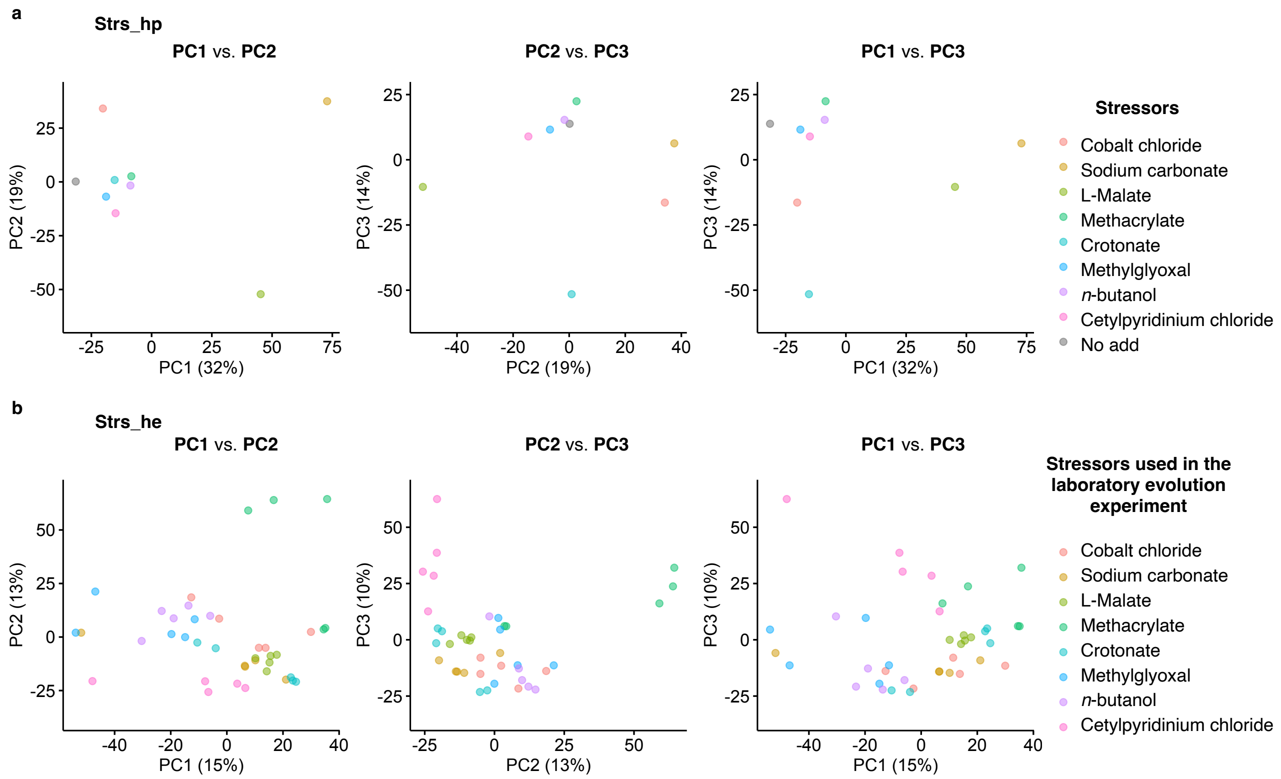


**Supplementary Fig. 21: Top three principal component (PC) scores for the Strs_hp and Strs_he datasets.**

Pairwise score plots for PC1–3 are shown for (**a**) Strs_hp and and (**b**) Strs_he. In panel **a**, the color legend indicates the stressors used to obtain the transcriptional profile of the ancestral strain. In panel **b**, the color legend indicates the stressors used in the laboratory evolution experiment (five independent lineages for each stressor). Transcriptome profiles of the isolates were obtained in the absence of stressors.

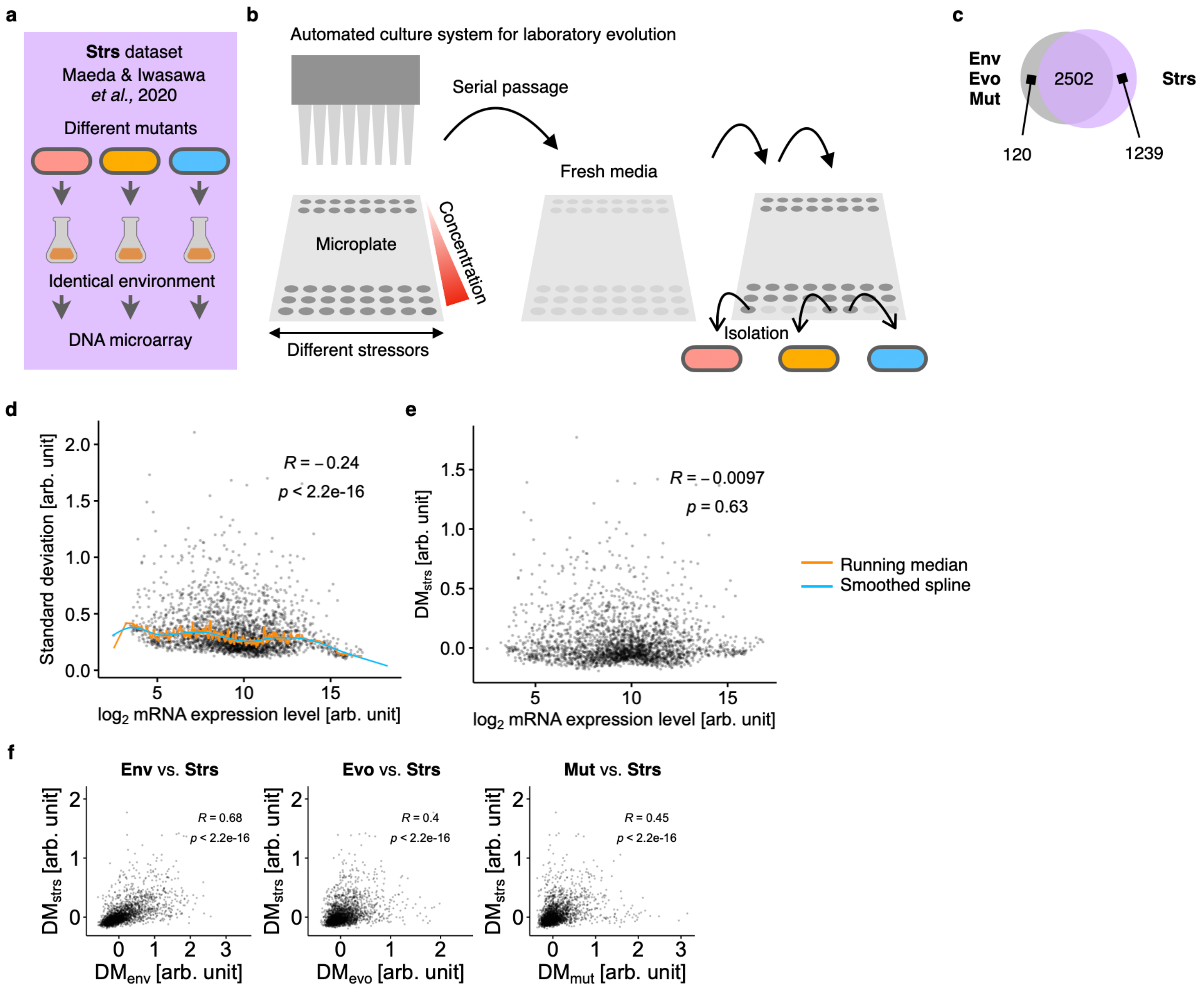


**Supplementary Fig. 22: The DM values in the Strs dataset.**

**a**,**b**, Schematic representation of the experiments conducted on the Strs dataset. These experiments were previously performed in our lab[6] using the same DNA microarray technique used in this study. **a**, In the Strs dataset, various mutants of *E. coli* MDS42 isolated from an evolutionary experiment under different stressors were subjected to DNA microarray analysis. Transcriptome profiles of the mutants were obtained under stress-free conditions. We analyzed the mutants that evolved in each of the 47 stressors (**Supplementary Data 10**) and in stressor-free conditions, excluding profiles from stressors used in the Env dataset. Thus, the Strs dataset reflects transcriptional variations induced by genetic perturbations through adaptive evolution. **b**, Evolution experiment to obtain mutants resistant to each stressor using MDS42 as the ancestral strain. The bacterial culture with the highest stressor concentration was selected every 24 h from cultures with varying stressor concentrations based on the defined cell density. The selected culture was serially transferred to a fresh medium with a stressor gradient. After 27 daily passages, mutants with improved resistance were isolated. **c**, Venn diagram illustrating shared genes among the datasets. 2502 genes were common. **d**, Relationship between mean expression levels and standard deviation. Each grey circle corresponds to a gene. The vertical deviation from a smoothed spline calculated from the running median of the standard deviations represents $DM_{strs}$. **e**, Relationship between the DM values and mean expression levels. **f**, Correlation of transcriptional variability between datasets. Pairwise correlations are shown for the DMs between Strs and the three main datasets (Env, Evo, and Mut) arranged from left to right. In panels **d**–**f**, Spearman's R and p-values (two-sided) are shown.

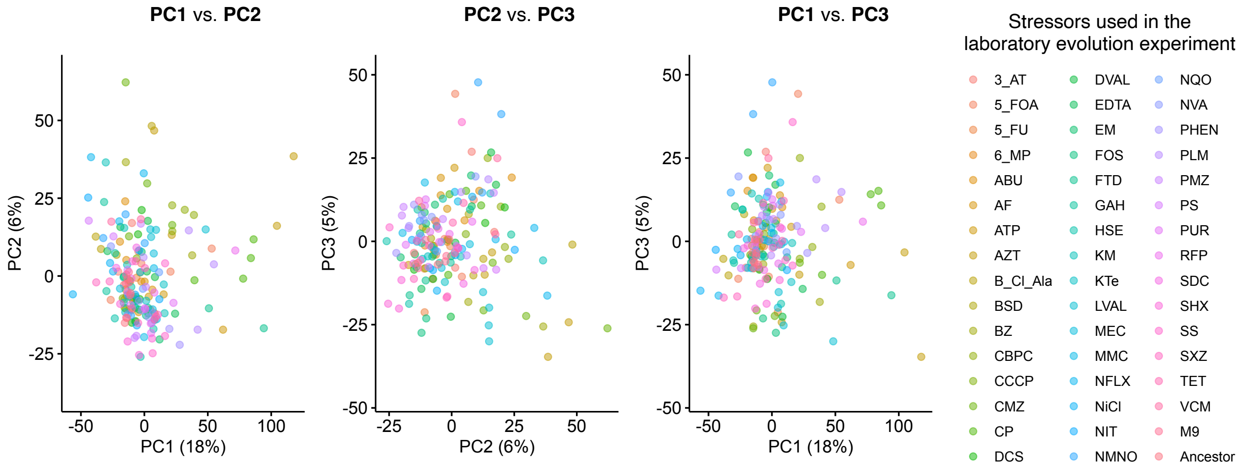


**Supplementary Fig. 23: Principal component analysis of the Strs dataset.**

Pairwise score plots of the top three principal components (PC1–3) in the Strs dataset are shown. The color legend indicates the stressors used in the laboratory evolution experiment, with four independent lineages for each stressor. Transcriptome profiles of the isolates were obtained in the absence of stressors. The label "M9" represents isolates from the stressor-free condition. Outlier profiles (only one out of 192 profiles) showing more than six standard deviations from the mean PC scores in any of the top three PCs were excluded from the analysis.

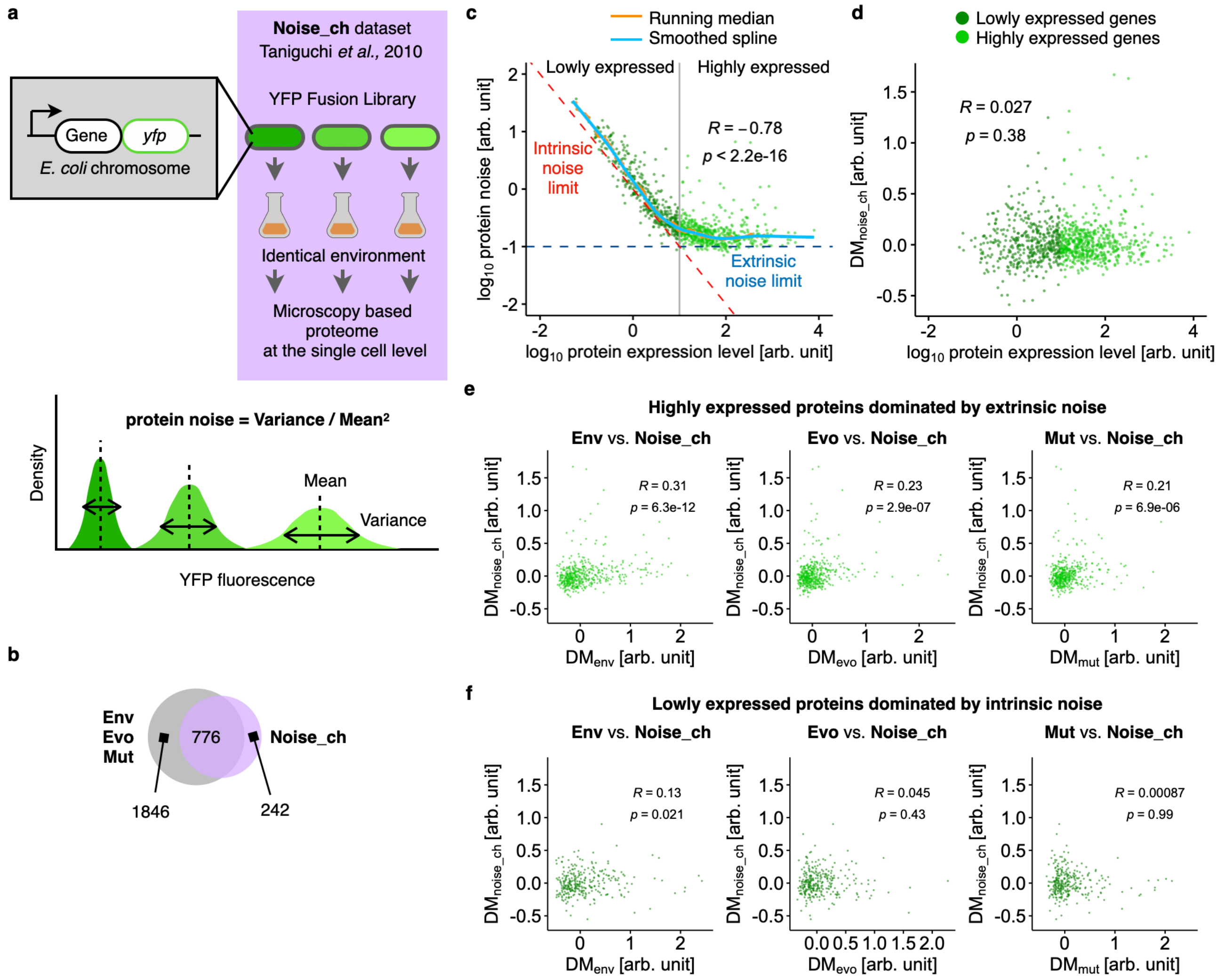


**Supplementary Fig. 24: The DM values in the Noise_chr dataset.**

**a**, Schematic representation of Noise_chr dataset. The dataset, as described by Taniguchi *et al.*[7], involved a library of yellow fluorescent protein (YFP) fusion strains of *E. coli* K12 cultured under identical conditions (top). Each chromosomal copy of the gene was labeled with YFP fusion, and YFP fluorescence was measured at the single-cell level by microscopy, resulting in the fluorescence distribution for each gene (bottom). Protein noise for each gene was defined as the variance divided by the square of the mean[7], representing isogenic cell-to-cell heterogeneity. **b**, Venn diagram displaying gene overlap between datasets. In total, 776 genes were identified. **c**, Relationship between the mean protein expression levels and protein noise. Each dot corresponds to a gene, and both axes were $\log_{10}$-transformed. The vertical deviation from a smoothed spline, calculated from the running median of the standard deviations, is denoted as $DM_{noise\_ch}$. The vertical line represents the expression level at which the intrinsic noise limit is equal to the extrinsic noise limit. Genes exhibiting lower expression levels than this line were expected to have a higher relative contribution of intrinsic noise to total noise, whereas genes with higher expression levels than this line had a higher relative contribution of extrinsic noise. This vertical line was used as a criterion to define lowly and highly expressed proteins. **d**, Relationship between $DM_{noise\_ch}$ and mean expression levels. **e**,**f**, Correlation of DM values between the datasets for highly (**e**) and lowly (**f**) expressed proteins. Pairwise correlations are shown for the DMs between Noise_ch and the three main datasets (Env, Evo, and Mut) arranged from left to right. Spearman's R and p-values (two-sided) are shown in the panels.

**Supplementary Note 1: Detailed descriptions of the five additional datasets**

**1. The Prtn dataset**

In the main text, we investigated whether transcriptional variability can extend to the protein level. For this purpose, we analyzed the proteomic variability of *E. coli* in response to various environmental conditions[4] (**Supplementary Fig. 17**). This dataset, referred to as the Prtn dataset, contains several environmental conditions similar to those used in the Env dataset, which was suitable for our purpose. Using this dataset, we calculated the distance of the standard deviation to a smoothed running median of the standard deviations of the $\log_2$-transformed protein abundance per cell across diverse environmental conditions. The resulting distance, $DM_{prtn}$, serves as a measure of proteomic variability in response to different environmental conditions (**Supplementary Fig. 17c,d**). Comparing DM values across different datasets, we observed that genes exhibiting higher DM values at the mRNA level tended to have higher $DM_{prtn}$ (Spearman's R = 0.27–0.43, **Supplementary Fig. 17e**). We also investigated whether the major axes of transcriptional variation were retained at the protein level. To this end, we performed PCA to identify the top three principal components (PC1–3) in the Prtn dataset (**Supplementary Fig. 18a**). The results of correlation analysis for the top three PCs between the Prtn and the three main datasets were provided in the main text (**Fig. 7c**). We observed that PC1 of the Prtn dataset was correlated with PC1 or PC2 in the Env, Evo, and Mut datasets. We also confirmed that PC1 in the Prtn dataset correlated with the growth rates at which the proteome profiles were obtained (Spearman's R=–0.96, **Supplementary Fig. 18b**), which was consistent with the results from the Env dataset (**Supplementary Fig. 8b**). Over all, the Prtn dataset provided protein-level variability in response to different environmental perturbations.

**2. The Strs_hp dataset**

In the main text, we investigated whether the magnitude and major directions of transcriptional variation in the three main dataset correlated with those in different environmental perturbations. For this purpose, we analyzed transcriptional profiles in response to environmental conditions that were not examined by the Env dataset. We used a compendium of transcriptional profiles obtained from a single strain of *E. coli*, MDS42, cultured under different stressors[5]. We removed the transcriptional profiles obtained under the same stressors as those in the Env dataset. The obtained dataset was called Strs_hp (**Supplementary Fig. 19**). Following the same procedure as for $DM_{env}$, we calculated $DM_{strs\_hp}$ as a measure of transcriptional variability in response to different environmental perturbations caused by stressors (**Supplementary Fig. 19e,f**). We observed strong positive correlations between $DM_{strs\_hp}$ and $DM_{env}$ (Spearman's R=0.62, **Fig. 7b**, **Supplementary Fig. 20a**) In addition, $DM_{strs\_hp}$ positively correlated with $DM_{evo}$ and $DM_{mut}$ (Spearman's R=0.36, 0.43, **Fig. 7b**). We also performed PCA to identify the major axes of transcriptional variation in Strs_hp (**Supplementary Fig. 21a**). The results of correlation analysis for the top three PCs between different datasets were provided in the main text (**Fig. 7c**). Compared to PC1–3 in the other datasets, we found that PC1 in the Strs_hp dataset correlated most strongly with PC1 in the Env dataset. Thus, the Strs_hp dataset provided transcriptional variability in response to different environmental changes that were absent in the Env dataset.

#### 3. The Strs_he and Strs dataset

In the main text, we also investigated whether the magnitude and major directions in the three main datasets were shared by transcriptional diversification accompanying adaptive evolution. Unlike neutral evolution driven by genetic drift, mutations accumulated in adaptive evolution constitute only a fraction of the mutations that are likely to be advantageous to an organism's fitness, depending on selective conditions. Accordingly, these biased mutations can affect the magnitude and direction of transcriptional diversification during adaptive evolution, which may violate the variation derived from the transcriptional variability identified in the main three datasets. To test this, we analyzed two datasets, Strs_he and Strs, comprising the transcriptome profiles of different mutants of *E. coli* isolated during adaptive laboratory evolution toward resistance to different stressors. The Strs_he dataset consisted of the transcriptome profiles of resistant mutants isolated from eight different stressful conditions[5] (**Supplementary Fig. 19**). The Strs dataset consisted of resistant mutants under 47 stress conditions[6] (**Supplementary Fig. 22**). The culture conditions under which the transcriptome profiles were obtained were stressor-free and identical between the mutants and the two datasets. Importantly, the transcriptional profiles derived from the mutants obtained under the same stress conditions as those in the Env dataset were removed. Following the same procedure as that for $DM_{mut}$, we calculated $DM_{strs\_he}$ and $DM_{strs}$ and used them as measures of transcriptional diversification through adaptive evolution. We found $DM_{env}$ strongly and positively correlated with $DM_{strs\_he}$ and $DM_{strs}$ (Spearman's R=0.56, 0.68; **Fig. 7b**, **Supplementary Figs. 20b, 22f**). Additionally, $DM_{strs\_he}$ and $DM_{strs}$ were moderately positively correlated with $DM_{evo}$ and $DM_{mut}$. We also performed PCA to identify the major axes of transcriptional variation in the two datasets (**Supplementary Figs. 21b**, **23**). Compared with PC1–3 in the other datasets, we found that PC1s in Strs_he and Strs correlated the most with PC1 in the Env dataset as explained in the main text (Spearman's R=0.48, 0.61; **Fig. 7c**). Overall, these two datasets provided transcriptional variation caused by genomic mutations accumulated through adaptive evolution.

#### 4. The Noise_ch dataset

Several studies have suggested that evolution tends to produce global canalization that buffers phenotypes against genetic and environmental perturbations[8]. Additionally, global canalization can buffer phenotypes against perturbations caused by stochastic protein noise[9, 10]. In the main text, we investigated this in *E. coli* using a dataset, termed the Noise_ch dataset, comprising cell-to-cell variations in the protein expression levels of genes encoded by endogenous promoters and natural chromosomal positions[7] (**Supplementary Fig. 24**). To compensate for the mean dependency of protein noise caused by intrinsic noise at lower expression levels[7], we followed a conventional method[11, 12] and calculated DM values as the vertical distance to a smoothed running median of $\log_{10}$-transformed protein noise, representing cell-to-cell variance divided by the square of the mean protein copy number per cell. We termed this relative protein noise $DM_{noise\_ch}$ a measure of variability against stochastic perturbations under identical environmental and isogenic conditions (**Supplementary Fig. 24c**,**d**). We found positive correlations (0.21–0.32, Spearman's R) between $DM_{noise\_ch}$ and the other DM values in

Env, Evo and Mut for highly expressed proteins (**Fig. 7b**, **Supplementary Fig. 24e**). In contrast, we observed only minimal or no significant positive correlations for lowly expressed proteins (**Supplementary Fig. 24f**), which is consistent with previous observations[13]. In contrast to a previous study, we strictly distinguished transcriptional variability in response to environmental perturbations from that in response to genetic perturbations. In addition, the protein noise we utilized was independent of plasmid copy number variation and potential confounding effects arising from gene expression from the plasmids. Despite these stringent considerations relative to the previous study, our results consistently revealed robust positive correlations between protein noise and transcriptional plasticity. Furthermore, our findings align with previous observations in yeast, where relative cell-to-cell variations in the protein expression levels of genes expressed from endogenous promoters and natural chromosomal positions correlated with transcriptional variability in response to genetic and environmental perturbations[12, 14]. Thus, the Noise_ch dataset provided protein-level variability in response to stochastic molecular noises.
